# Single seed microbiota: assembly and transmission from parent plant to seedling

**DOI:** 10.1101/2021.05.31.446402

**Authors:** Guillaume Chesneau, Béatrice Laroche, Anne Préveaux, Coralie Marais, Martial Briand, Brice Marolleau, Marie Simonin, Matthieu Barret

## Abstract

Gaining basic understanding of processes involved in seed microbiota assembly is a prerequisite for improving crop establishment. Investigation of microbiota structure during seed development revealed that individual seeds of bean and radish were associated with a dominant bacterial taxon representing more than 75% of all reads. The identity of this taxon was highly variable between plants and within seeds of the same plant. Succession of dominant taxa occurred during seed filling and maturation through *Selection*. In a second step, we evaluated seed to seedling transmission of these dominant seed-borne taxa. We showed that initial bacterial abundance on seeds was not a good predictor of seedling transmission and that the identity of seed-borne taxa can impact seedling phenotype. Altogether this work unveiled that seeds are colonized by few bacterial taxa of highly variable identity, which appears to be important for the early stages of plant development.

## Introduction

Seed microbiota allows the transmission of microbial communities between plant generations^1^ and is a key factor influencing both seed vigor and seedling development, two essential steps for crop establishment. Several studies focused on the role of the mature seed microbiota in releasing dormancy^2^, improving germination^3^ or protecting against damping-off^4^. In addition, seed microbiota may have longer-term consequences on crop establishment by promoting establishment of soil-borne plant-beneficial microorganisms^5^ or limiting the incidence of soil-borne plant pathogens^6^. However, all the above-mentioned studies characterized microbiota of dried mature seeds, while limited knowledge is available on the dynamics of microbiota assembly during seed development^7^. A consequence of this knowledge gap is that the origin, timing of arrival and succession of seed-associated microbial taxa is mostly unknown, as well as their future consequences on seed vigor and seedling development.

To date, the characterization of seed microbiota structure has been performed on about 50 plant species including major crops such as barley ^8^, bean^9^, rapeseed^10^, rice^11^, tomato ^12^ and wheat^8^. *Selection* by the local environment or by the host significantly modulates seed microbiota composition^9, 10, 13^. Other ecological processes such as dispersal by pollinators^14^ and ecological drift^15^ are also important drivers of seed community assembly. However, in most studies seed microbiota was characterized on large seed samples from 10 to 1,000 seeds. One reason for using seed lots rather than individual seeds is related to the technical difficulties of collecting enough microbial DNA from a single seed as these are usually colonized by a low bacterial population size^16^. Another rationale for using seed samples is related to historical studies carried out in seed phytopathology. Indeed, seed-transmitted plant pathogens are generally distributed within a Poisson distribution^17^ and increasing sampling size increases the probability of detection. While working on seed lots is valid for improving plant pathogen detection, this sampling procedure artificially increases seed microbiota richness and more importantly does not allow the estimation of seed-to-seed variability.

The few studies carried out at the individual seed level or small seed samples (i.e. less than 10 seeds) through culture-dependent^16, 18^ or –independent^19–21^ approaches have revealed a low microbial richness. It has even been proposed that each seed was colonized by none or only one endophytic microorganism^18^. The authors of the latter study propose a hypothesis, the “primary symbiont hypothesis”, that endophyte microbial transmission in seed is bottlenecked as a result of plant defenses and interactions among seed-transmitted microorganisms^18^. However, this hypothesis has still not been confirmed at the whole individual seed microbiota level, which includes endophyte and epiphytes. In addition, the order and routes of arrival of these “primary symbionts” as well as the ecological processes involved in their assembly remained unexplored.

The presence of a microorganism, on, or in seeds is not a guarantee of transmission to the seedling^22^. To date studies focusing on the dynamics of microbial communities during germination and emergence have been performed on several subsamples of the same seed lot (e.g: ^8, 12, 23–25)^. However, in order to truly assess transmission of seed-borne bacterial taxa to the seedling, it is necessary to work at the individual seed-seedling level in a non-destructive way.

In this context, this study aimed to address several fundamental questions on the dynamics of seed microbiota from mother plant to seedling. Firstly, does the order and timing of microbial taxa immigration during seed development impact seed microbiota composition? In other words, are the first microorganisms that colonize the seed during its development, the ones that will be established on mature seeds? What are the main seed transmission pathways employed by these seed-borne taxa? What are the ecological processes involved in seed microbiota assembly? What is the dynamics of seed-borne taxa during seedling emergence? And what is their impact on seed germination and seedling phenotype?

To answer these questions we worked on seeds of common bean (*Phaseolus vulgaris*) and radish (*Raphanus sativus*), two plant species whose seed microbiota has been extensively characterized ^7, 9, 15, 23^. We estimated microbiota composition of single seeds of different individual plants during seed filling and maturation through amplicon sequencing. We also inferred the origins of seed-associated taxa through sampling and sequencing of flowers, vascular tissues and atmosphere microbiota. In a second step, we evaluated seedling transmission of seed-borne bacteria by sampling, in a non-destructive way, individual seed and seedling microbiota. This second step was performed aseptically in order to specifically focus on the inheritance of the seed microbiota, avoiding influence of the environment. Finally, we assessed the impact of seed microbiota on seedling emergence and confirmed the impact of some selected taxa on seedling fitness. We showed that individual seeds are colonized by one dominant taxon with variable taxonomic identity. These taxa are not necessarily abundant in the future seedling but can have a significant beneficial or detrimental impact on the seedling phenotype, such as *Pantoea agglomerans* on bean seedling.

## Results

### Experiment 1 – Culture-based enrichment of seed-borne bacterial strains efficiently captured the diversity of the seed microbiota

Sequencing of samples with low microbial biomass is subject to a low signal to noise ratio as a result of weak amounts of DNA starting material^26^. We assessed whether culture-based enrichment of seed-borne bacteria could increase population size without distorting community structure. Firstly, the bacterial diversity obtained after dilution and plating on nine culture media was estimated through amplicon sequencing of two molecular markers that differed in their taxonomic resolution, namely the V4 region of 16S rRNA gene (hereafter referred to as 16S) and a portion of *gyrB*^23^. According to Faith’s phylogenetic diversity, 1/10 strength Tryptic Soy Agar (hereafter referred to as TSA10) allowed the greatest diversity of bacterial isolates to be obtained (**Fig. S1**), therefore confirming previous results obtained with root-associated bacterial strains^27^.

In a second step we compared the bacterial community profiles before and after culture-enrichment on TSA10 on 24 bulk seed samples of bean and radish. In total 1,276 ASVs were detected with 45 common to both approaches (**Fig. 1A**) including the most abundant ASVs (**Fig. 1B**). ASVs specifically detected with one of the two approaches were not affiliated to a particular phylogenetic lineage (**Fig. 1C**). Hence, differences in the detection of taxa with or without TSA10 culture-enrichment could be due to differences in amplification of rare taxa. Altogether, these results highlighted that investigating bacterial community assembly is possible with a step of culture-based enrichment and may even allow better detection of rare taxa since 927 ASVs were specifically detected following culture-based enrichment (**Fig. 1A**).

**Figure 1:**
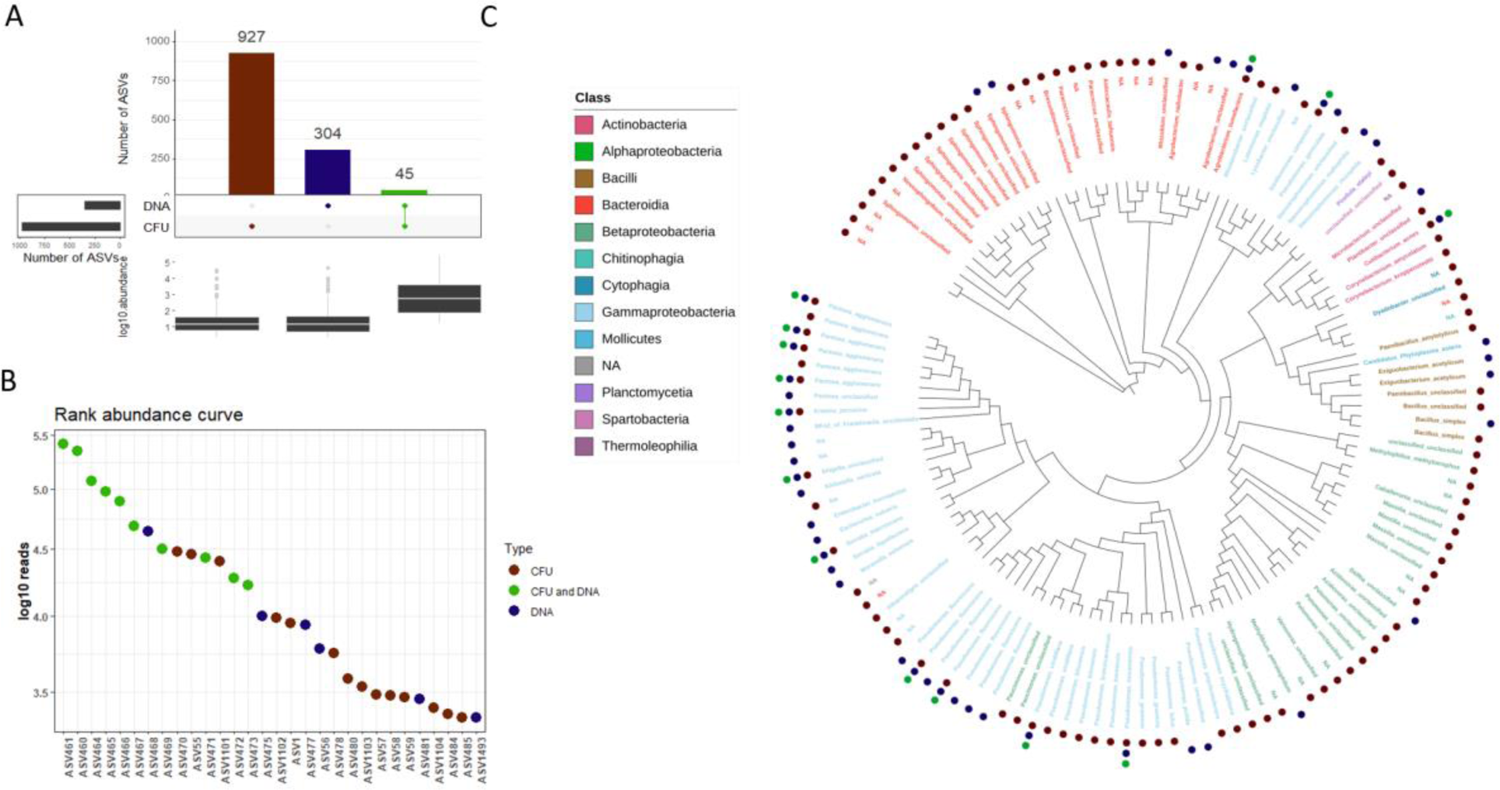
Comparison of seed bacterial community profiles before and after culture-enrichment on TSA10. **A.** Visualization of the number of *gyrB* ASVs specifically detected before (DNA) and after (CFU) culture-enrichment, or jointly (CFU and DNA). Log10 abundance of ASVs are displayed in the boxplots. **B.** Rank abundance curve of the 30 most abundant ASVs. **C.** Neighbor-Joining phylogenetic tree of the most abundant ASVs (minimum relative abundance of 1%). Branch tips colors correspond to the Class taxonomic level. Blue and red circles represented ASVs specifically associated with culture-independent and -dependent approaches, respectively. Green circles represented ASVs detected with both approaches.

### Experiment 2 – Assembly and structure of the microbiota during individual seed development

#### Percentage of seeds colonized by cultivable bacteria varied between plants

Seeds from bean and radish were collected aseptically during seed filling and maturation, two stages defined by measuring seed water content (**Fig. 2A**). As expected the percentage of water content significantly decreased during seed development for both plant species, especially during late maturation. Bacteria were detected on average in 43% to 84% of bean seeds per stage and 30% to 82% of radish seeds per stage (**Fig. 3A**). Variation in the percentage of seed-containing bacteria was associated with the sampling stage for both plant species (*P*<0.01). While no correlation between bean seed development and percentage of seed-containing bacteria was observed, the percentage of seed-containing bacteria gradually increased during radish seed development (**Fig. 3A**). Within each sampling stage, the percentage of detection of seed-borne bacteria was significantly (*P*<0.01) different between individual plants and varied from 25% to 100% per bean plant and 20% to 100% for radish (**Fig. 3A**).

**Figure 2:**
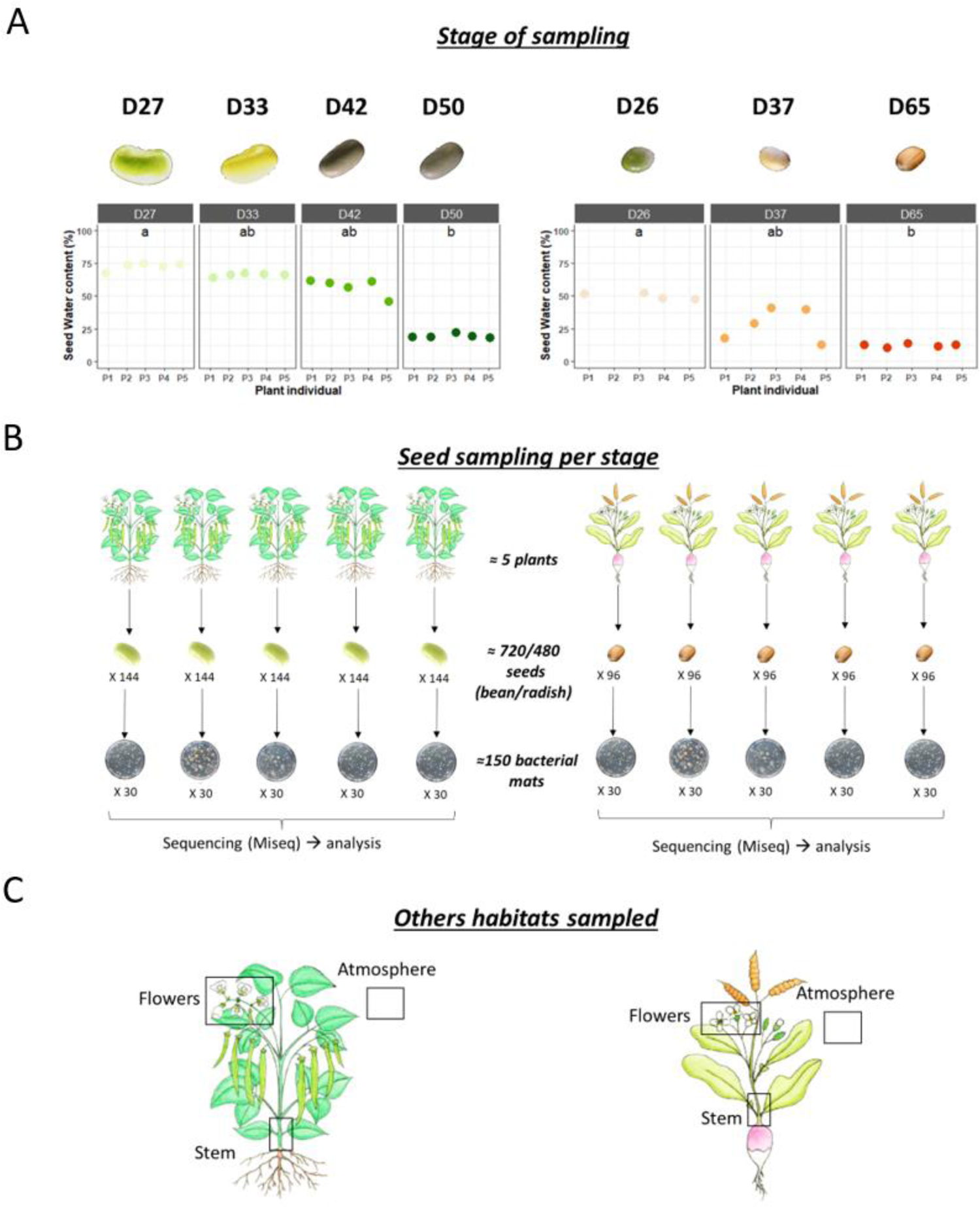
Experimental design used to monitor the assembly of bacterial communities during seed development. **A.** Seed water content (%) associated with samples collected during bean (left) and radish (right) seed development. Letters denote significance evaluated from Kruskall-Wallis following Dunn test *post hoc* (*P* < 0.01) **B.** Number of samples processed at each stage. **C.** Sampling of flowers, stems and atmosphere, which could represent the main inoculum sources for seed taxa.

**Figure 3:**
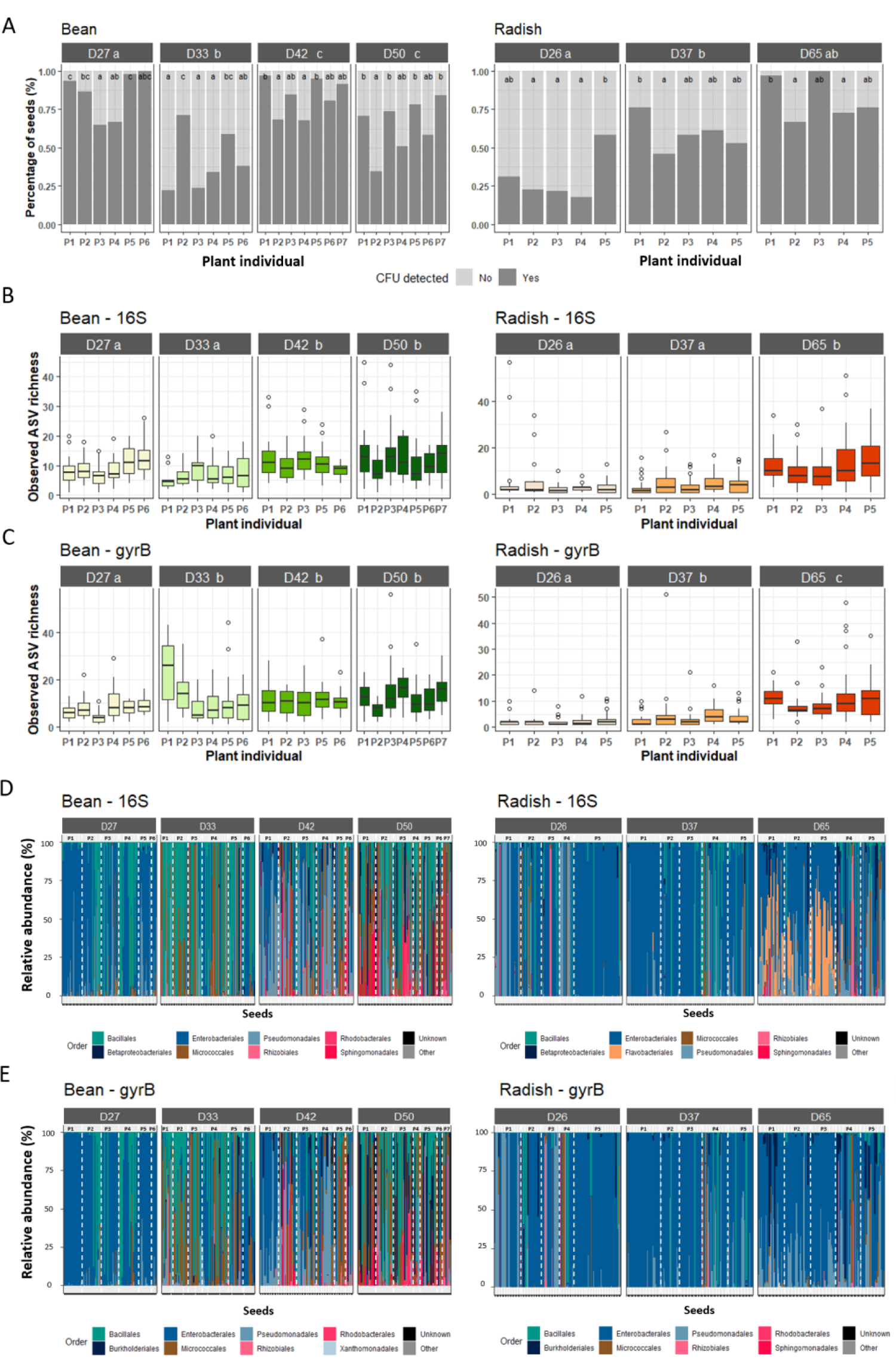
Structure of the bacterial fraction of the seed microbiota. **A.** Percentage of seeds with (dark grey) or without (light grey) CFU detected. **B-C.** Number of ASV per seed (richness) estimated with 16S (B) or *gyrB*. Seeds were grouped according to their plant of origin (x-axis). **D-E.** Relative abundance of bacterial Orders for each individual seed according to 16S (**D**) or *gyrB* (**E**). White dotted lines represented individual plants. The names in each frame correspond to the stages sampled during seed development. The letters (a, b, c) indicated the significance level (*P* < 0.01).

#### Individual seeds are associated with low bacterial richness

Few bacterial taxa were detected per individual seeds. Indeed, a median richness of eight 16S ASVs in bean and four 16S ASVs in radish were observed (**Fig. 3B**), while a median of nine and three *gyrB* ASVs were observed in bean and radish seeds (**Fig. 3C**). Bacterial richness significantly (*P*<0.01) increased during seed development of both plant species regardless of the molecular markers employed. Moreover, within each sampling stage, high variability in richness between seeds was observed (**Fig. 3B and 3C**). For instance, based on 16S, one to 45 ASVs were detected per bean seed at the mature stage (D50, **Fig. 3B**), while one to 51 ASVs were associated with radish seeds at the same developmental stage (D65, **Fig. 3B**). Variation in richness was accompanied by a high variation in taxonomic composition (**Fig. 3D and 3E**). Changes in phylogenetic composition between seed bacterial communities was significantly (*P*<0.001) explained by the seed development stages for bean (16S: 18.3%; *gyrB*: 16.1%,) and radish (16S: 15.3%; *gyrB*: 7.4%). In addition, variation in phylogenetic composition was also significantly (*P*<0.001) associated with plant inter-individual variations in bean (16S: 26.5%; *gyrB*: 26.1%) and radish (16S: 28.5%; *gyrB*: 20.0%).

#### A seed is associated with one dominant bacterial taxa

One dominant bacterial taxa was associated with bean and radish seeds. According to 16S (**Fig. 4A**), these dominant taxa had mean relative abundance of 77.1% (± 22.0%) and 75.4% (± 22.8%) with *gyrB* (**Fig. S2A)**. The identity of the dominant taxa was highly variable between seeds of both plant species with 94 16S ASVs and 285 *gyrB* ASVs distributed in more than 15 bacterial orders (**Fig. S3** and **S4**).

**Figure 4:**
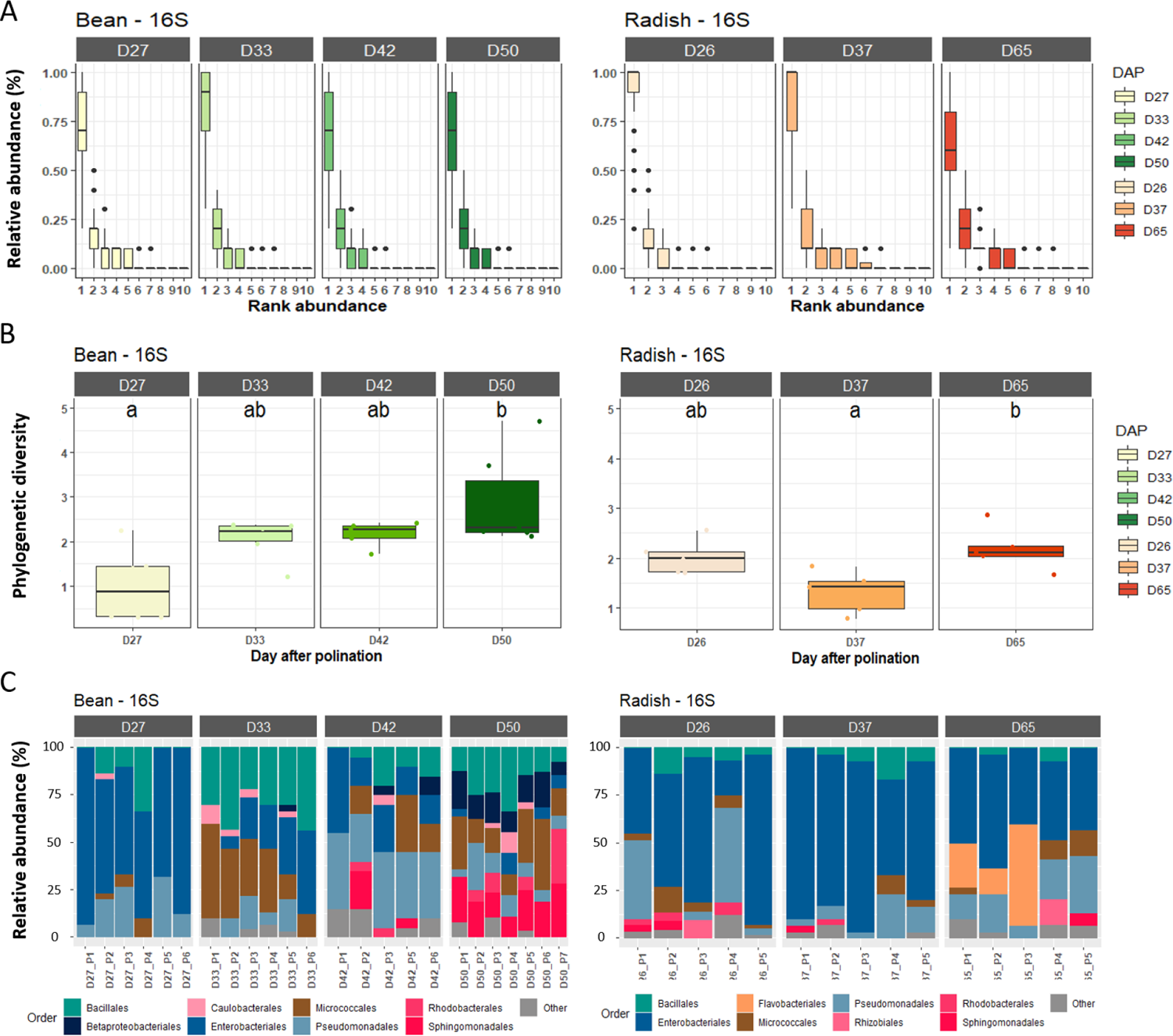
Dominant bacterial taxa within the seed microbiota based on 16S. **A.** Rank abundance curves of bacterial ASVs associated with individual seeds. **B.** Faith’s phylogenetic diversity of rank1 ASV agglomerated at the plant level. **C.** Relative abundance of bacterial Orders calculated from rank 1 ASV agglomerated at the plant level. The names in each frame correspond to the stages sampled during seed development. The letters (a, b, c) indicated the significance level (*P* < 0.01).

The phylogenetic diversity of dominant taxa per plant significantly (*P*<0.05) increased during bean seed development (**Fig. 4B**). Moreover, a clear shift in bacterial taxonomic composition was observed during bean seed development with a decrease in the occurrence of dominant taxa affiliated to Enterobacterales (**Fig. 4C** and **S2C**). Although our destructive sampling does not allow a direct monitoring of microbiota dynamics during seed development, phylogenetic diversity and taxonomic composition indicated succession of dominant taxa over seed filling and maturation of bean. For radish, we did not observed significant change in phylogenetic diversity. However, changes in taxonomic composition of dominant taxa per plant species was observed with an increase in occurrence of Flavobacteriales during the last seed maturation stage (**Fig. 4C**).

Within a sampling stage, a large variation in taxonomic composition was observed between seed bacterial communities from different plants (**Fig. 3D**). The distribution pattern of seed dominant taxa indicated a significant (Chi-squared test, *P*<0.01) association between some dominant taxa and specific plant individuals (**Fig. S3**, **S4** and **Table S1**). For instance at D27 in bean, one dominant taxa affiliated to Erwiniaceae with 16S and *gyrB* was significantly more frequent in plant 1 (**Fig. S3**, **S4** and **Table S1**). Hence the identity of dominant seed-associated taxa appeared to be dependent on the local pool of bacterial taxa found within one specific plant individual.

#### The origin of seed-associated taxa

To gain more insight on the potential inoculum sources of the seed microbiota, we performed additional sampling of the bacterial fraction of the microbiota after culture enrichment on three distinct habitats, namely flowers, stems and atmosphere (**Fig. 2C**). These three habitats could represent the main sources of the floral, internal and external pathways described for seed transmission^28^. Detection of *gyrB* sequence variants was employed in this section as *gyrB* has a better taxonomic resolution (i.e. infraspecific) than 16S.

Observed richness associated with atmosphere, flowers and stems was lower than on seeds for most of the seed developmental stages (**Fig. S5A**). All communities were dominated by bacteria belonging to the Proteobacteria, Actinobacteria and Firmicutes (**Fig. S5B**). The temporal dynamics of the airborne samples over time was explaining 33.7% (*P* < 0.001) of variability in phylogenetic composition (**Fig. S5C)**. We also observed significant (*P* < 0.05) variation in phylogenetic composition of bacterial communities associated with bean stem (23.0%) and radish stem (21.4%) over time (**Fig. S5C)**. The percentage of seed taxa that were not detected in atmosphere, flowers and stems was at least higher than 80% for both plant species at each sampling time (**Fig. 5**). Overall, we detected more seed-associated ASVs in the atmosphere compared to the other habitats sampled (**Fig. 5**).

**Figure 5:**
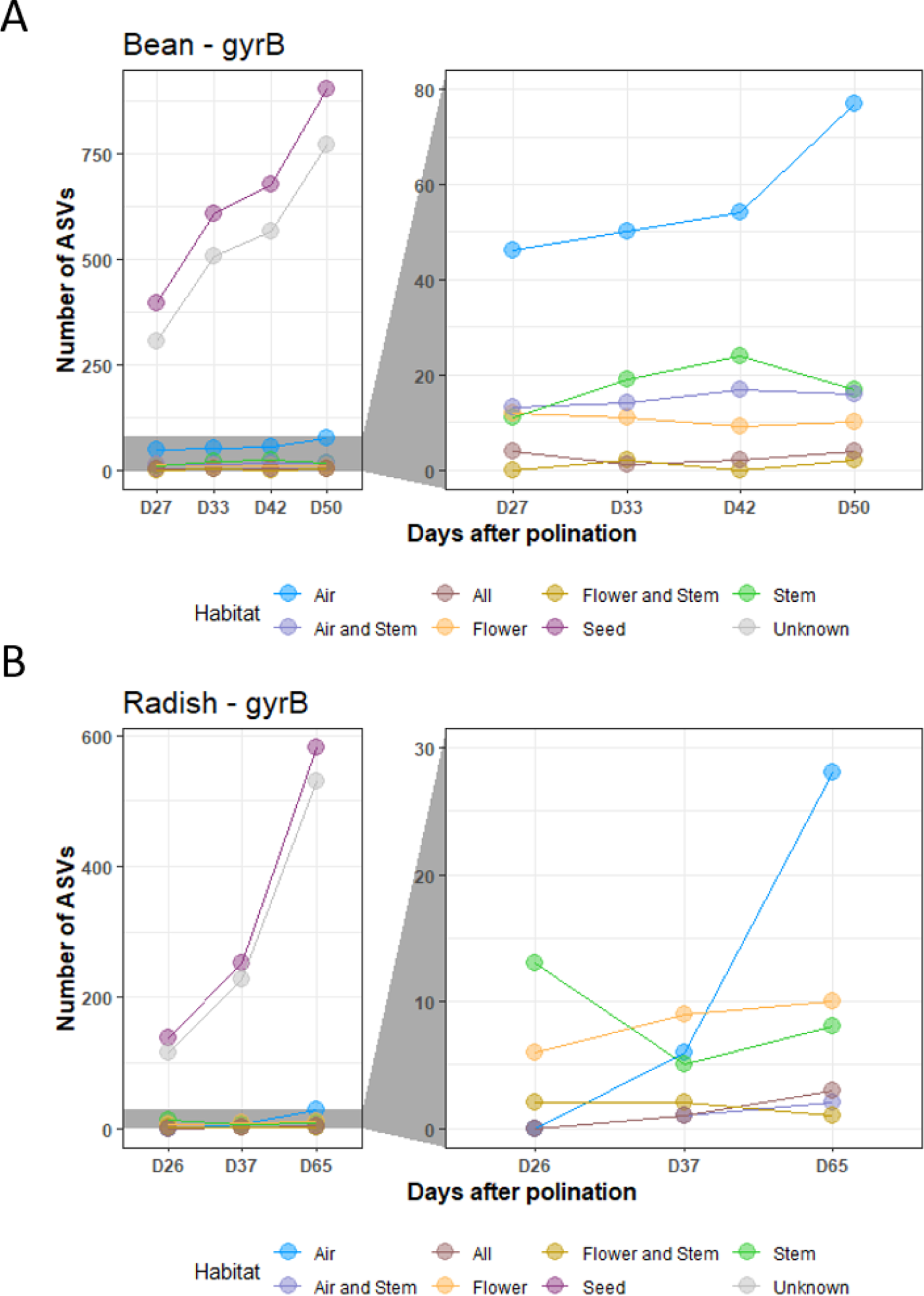
Origin of seed-borne taxa. **A-B.** Number of *gyrB* ASVs associated with bean (**A**) and radish (**B**) seeds that were detected in other sampled habitats: flowers, stems and atmosphere. The right panel is a zoom of the left panel.

We next looked at the origin of the dominant taxa. The most frequent dominant ASV, *Pantoea agglomerans* (30% of all seeds with 289 occurrences, **Fig. S4**), was detected in all sampled habitats for both plant species (**Fig. S6**). Although some taxa specifically associated with one specific plant (**Table S1**) were highly abundant in stem (e.g. ASV5), this observation was not generalized to the whole, therefore suggesting distinct transmission routes (**Fig. S6**).

#### Dispersion is less important than local seed processes

Since dominant seed-associated taxa were specifically associated with one plant individual, proportions of observed ASVs within each seed were aggregated at the plant level to form a species-proportion distribution. We compared the resulting distribution with the observed species proportions predicted by an adapted neutral model (see Methods). Species-proportion distributions be considered as a first order approximation^29^ for assessing the relative importance of dispersal over seed local processes (i.e. ecological drift or *Selection*) through the ratio *I* as well as the size *S* of the species pool available for these processes (see Methods).

Dispersion was less important than local seed processes in more than 85% of bean and radish plants. For bean, no clear pattern appeared over time but *I* was less important at D33 than at the following stages (**Fig. 6A and 6C**, *P*=0.003 for 16S, *P=*0.063 for *gyrB*). For radish, *I* increased significantly (*P=*0.012 for 16S, *P=*0.05 for *gyrB*) between D37 and D65 (**Fig. 6E and 6G**). The species pool size available for these processes was markedly low with median values ranging from 3-6 and 2-5 for bean and 2-6 and 3-4 for radish, for *gyrB and* 16S markers respectively (**Fig. 6B, 6D, 6F and 6H**). These results suggest that the balance between dispersion and local processes is shifted in time.

**Figure 6:**
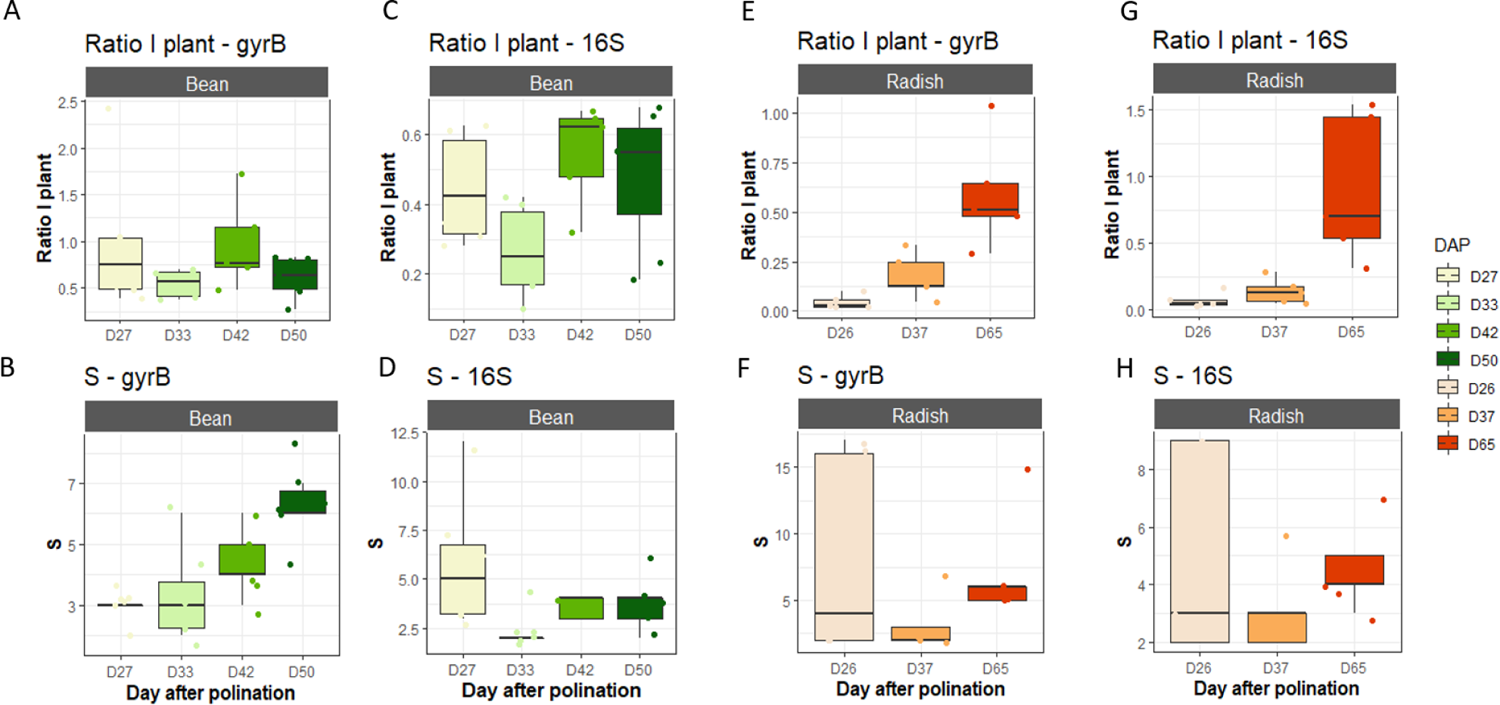
Relative importance of dispersal over seed internal processes. **A-C-E-F.** The ratio *I* is used to estimate the relative importance of dispersal over seed internal processes (see Methods). **B-D-F-H.** Distribution of S (richness) for bean plants (green) and radish plants (red) as a function of the sampling day. Data were estimated with *gyrB* (**A-B-E-F**) and 16S (**C-D-G-H**).

#### Selection drive local seed microbiota assembly at the plant level

To estimate the relative importance of *Selection* and ecological drift in the local (e.g. plant level) assembly of the seed microbiota we employed null model or more specifically abundance-based β-null deviation measures ^30^. With respect to the data obtained on bean seeds with 16S and *gyrB*, β-nearest taxon index (βNTI) values negatively deviated from the null expectation (**Fig. 7A** and **B**). βNTI values less < −2 are considered as significantly different from the null expectation ^30^. Since all βNTI values were below this threshold, this indicated that the phylogenetic composition of bean seed communities aggregated at the plant level are more similar than expected by chance and that this similarity could be driven by *Selection* ^31^.

**Figure 7:**
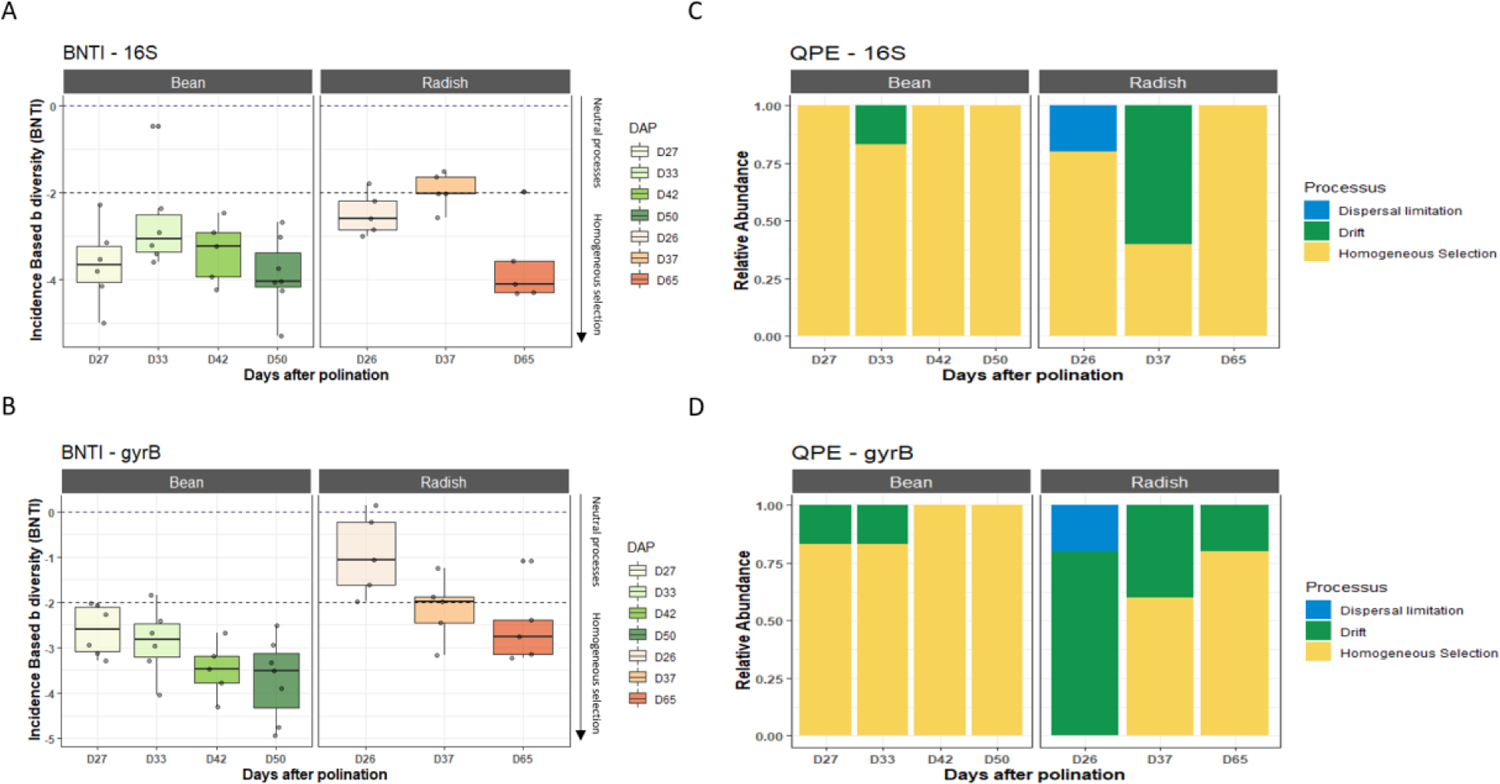
Ecological processes involved in seed microbiota assembly at the plant level. **A-B.** β-nearest taxon index (βNTI) values were calculated with 16S (**A**) and *gyrB* (**B**) to assess deviation from the null expectation. **C-D.** Relative importance of species-sorting (homogeneous selection), dispersal limitation or ecological drift in assembly of seed bacterial communities at the plant level. Analyses were performed with 16S (**C**) and *gyrB* (**D**).

Regarding the bacterial communities associated with radish seeds, βNTI values were distributed around −2 with 16S and *gyrB* for the two first sampling stages and then less than −2 at the final mature seed stage (**Fig. 7A** and **B**). Therefore, it seems that *Selection* had less influence on assembly of radish seed communities, at least on the early seed developmental stages, in comparison to bean seed communities. Thanks to quantitative process estimates (QPE) framework^30^, we were able to estimate the relative importance of other ecological processes during assembly of the seed microbiota. Based on this framework, it seems that ecological drift and dispersal limitation occurred during the early assembly stages (D26 and D37) of radish seed microbiota (**Fig. 7C** and **D**). At the late seed maturation stage of radish (D65), *Selection* became the principal process of community assembly (**Fig. 7C** and **D**).

### Experiment 3 – Seedling transmission of seed-borne taxa and their influence on seed vigour

#### Seed microbiota composition is influenced by leaf height

In the previous section, we showed that individual seeds were colonized by a few bacterial taxa of variable identity. Part of the variation was explained by the individual plants. To gain more insight into the factors involved in this variability at the seed level, mature seeds from five individual plants were collected from eight fruits located at four different leaf heights for each plant species. Based on nested PERMANOVA performed on weighted UniFrac distance, leaf height inside each plant explained 12% and 11.8% of variation (*P* <0.05, **Fig. S7A** and **S8A**) in bacterial phylogenetic composition of bean and radish seeds, respectively. In contrast, we did not observe a significant pattern of richness and taxonomy according to the position of the seed according to the leaf height (**Fig. S7B** and **S8B**). We confirm here the hypothesis of a spatial heterogeneity of seed microbiota between plants but also within each plant individual.

#### Dynamics of seed microbiota during seedling emergence

We next characterized which seed-borne bacterial taxa were transmitted to the seedling. Dynamics of the seed microbiota during emergence were analyzed in soil-less conditions using a non-destructive sampling procedure (see Methods), which makes it possible to monitor these dynamics at the level of individual seedling. We analyzed only the samples where CFU were recovered for seed and seedlings, therefore excluding non-germinating seeds or seeds without detectable CFU (**Fig. 8A**).

**Figure 8:**
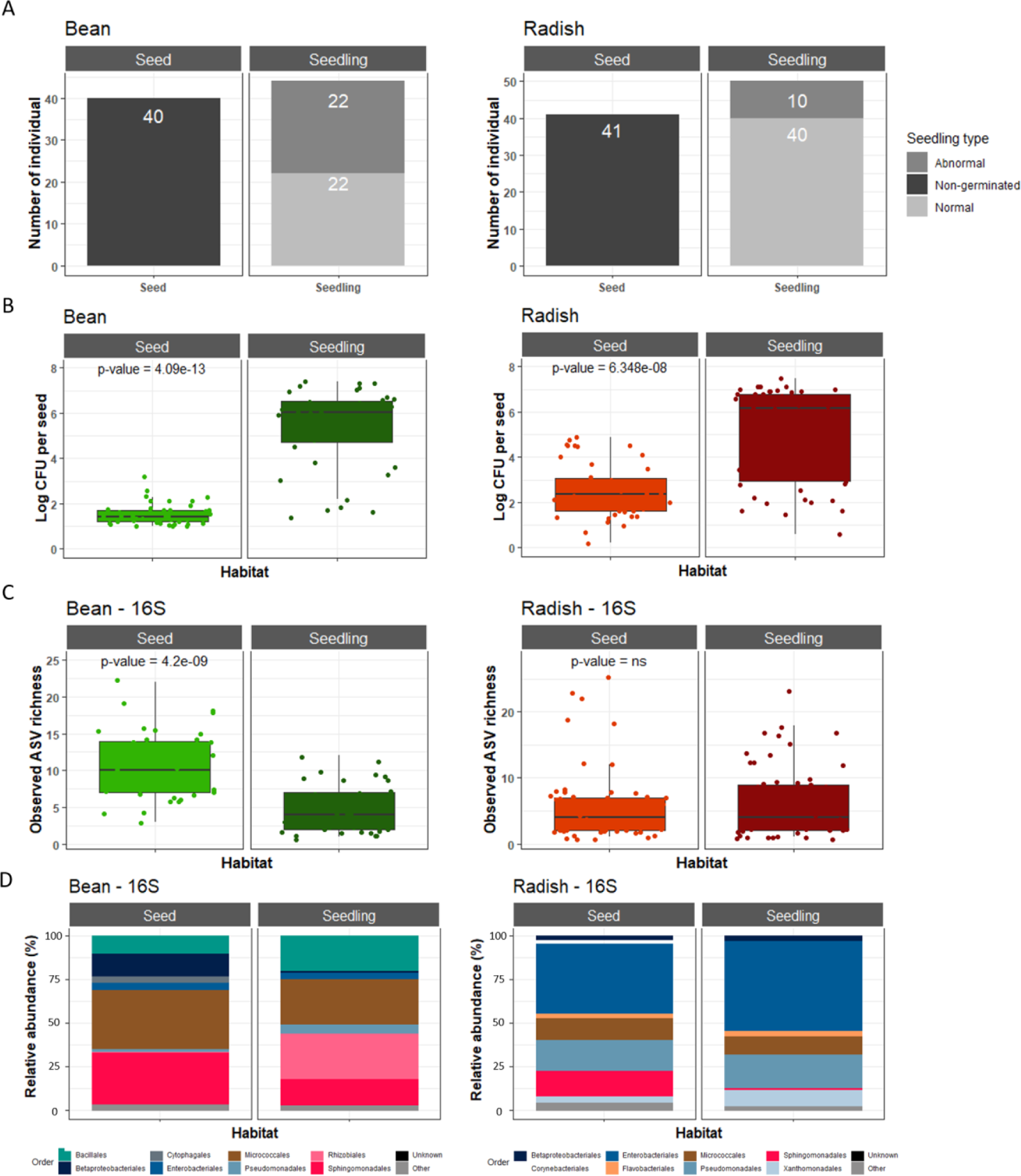
Comparison of seed and seedling microbiota profiles. **A.** Number of non-germinated seeds (dark grey), abnormal (grey) and normal seedling (light grey). **B.** Average bacterial population size (log10 CFU) associated with seed and seedling. **C.** Average richness (estimated with 16S ASVs) in seed and seedling. **D.** Taxonomic profile (Order level - 16S) of seeds and seedlings microbiota. *P*-values were derived from a Wilcoxon rank sum test.

During transition from seed to seedling, median bacterial population size increased by 1.4 to 6.0 log10 CFU in bean and by 2.3 to 6.1 log10 CFU in radish (**Fig. 8B**). Increase in bacterial population size during emergence was associated with a significant (*P*<0.001) decrease in richness in bean (10 to 4 ASVs with 16S and *gyrB*). In contrast, no significant change in bacterial richness was detected in radish with both molecular markers (**Fig. 8C** and **S9**).

About one fifth (43 out of 223) and one quarter (56 out of 243) of the bacterial ASVs associated with bean and radish seeds were detected in seedlings using 16S (**Fig. 9A**). The use of *gyrB* reduced these proportions to 11% and 5.6% for bean and radish, respectively (**Fig. S10A**). Although seed-borne ASVs not detected in the seedlings were affiliated with a multitude of bacterial orders, Burkholderiales were almost never detected in seedlings of both plant species (**Fig. S11** and **S12**). More surprisingly some seedling-associated ASVs were not detected in seeds. The absence of CFU detection in our sterile experimental system (**Fig. S9A**) strongly suggested that these seedling-associated ASVs originated from seeds. Among these ASVs specifically associated with seedlings were numerous Bacillales and Rhizobiales for bean and multiple Pseudomonadales for radish (**Fig. S11** and **S12**).

**Figure 9:**
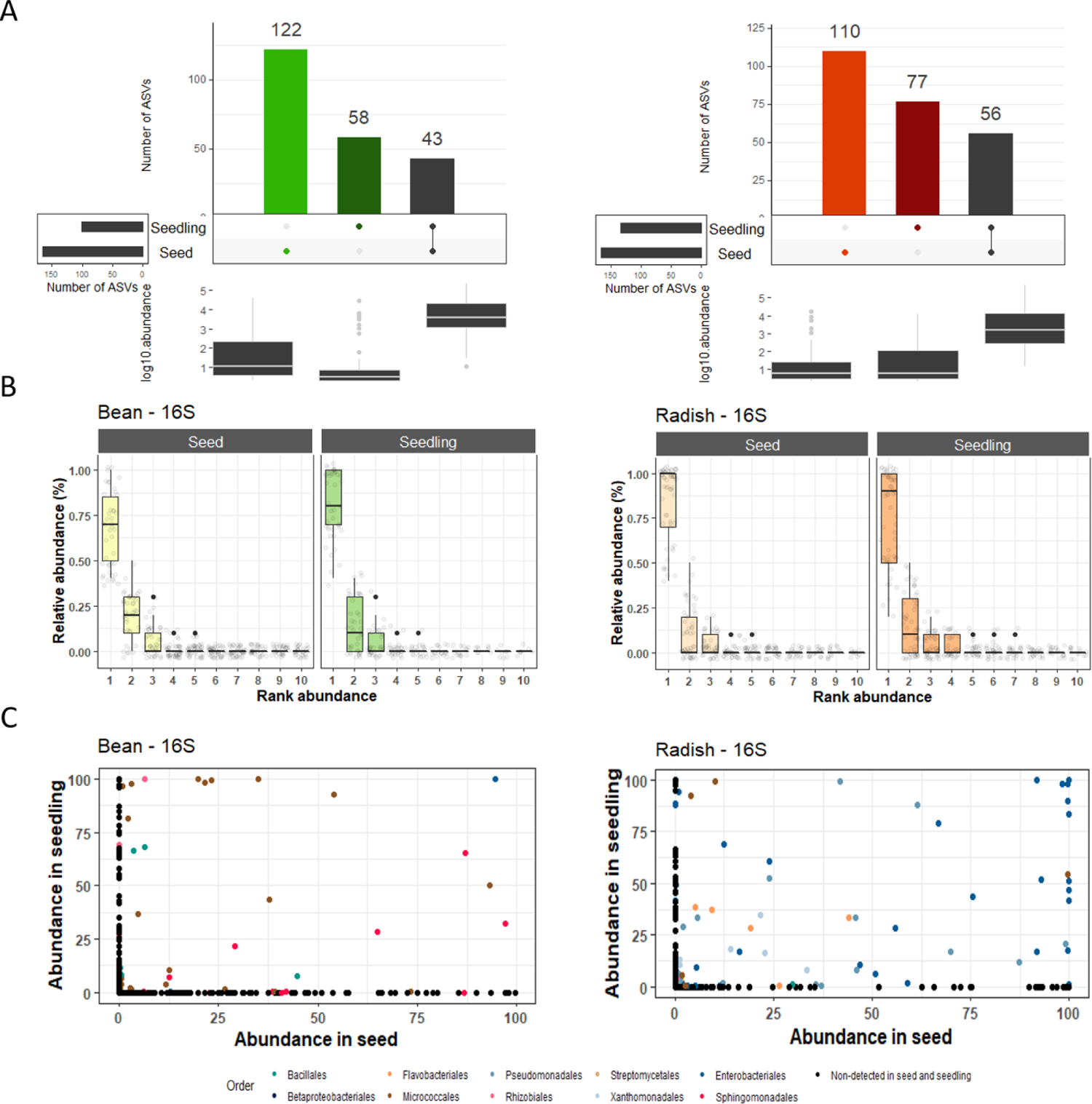
Seed to seedling transmission. **A.** Number of 16S ASVs specifically detected on seeds (light color), seedlings (dark color) or both habitats (black). **B.** Rank abundance curves of 16S ASVs in individual seeds and seedlings. **C.** Comparison of 16S ASV relative abundance in seed and seedling. Each dot corresponded to one ASV, which is colored according to its taxonomic affiliation at the Order level.

#### The initial abundance of seed-associated taxa does not explain its transmission to seedlings

Since transmission of some seed-borne plant pathogens to seedling is explained by their initial populations sizes on seeds ^32, 33^. We investigated whether the relative abundance of seed-borne ASVs was a predictor of seedling transmission. As what had already been observed on experiment 2, each individual seed of both plant species was associated with one dominant ASV. The same pattern was observed for seedling, where one dominant ASV was associated with bean and radish seedlings (**Fig. 9B** and **S11B**).

However, the dominant seed-borne ASV did not necessarily become dominant in the corresponding seedlings. Indeed, only 11 (out of 57) and six (out of 52) dominant seed-borne ASVs were also dominant on radish seedling according to 16S and *gyrB*, respectively. This value is even lower for bean with four 16S ASVs (out of 43) and one *gyrB* ASV, which were dominant in both habitats. Altogether these results suggest that the initial abundance of seed-borne taxa is not predictive of its transmission to seedlings as many rare seed-borne taxa become dominant on seedlings, especially in bean.

#### The identity of the seed-borne taxa affected seedling fitness

Since seeds are colonized by few taxa of variable identity, we investigated whether the identity of these ASVs could impact the first phases of crop establishment, namely germination and seedling emergence.

A high proportion of seeds (48% for bean, 45% for radish) did not germinate under our experimental conditions (**Fig. 8A**). Bacterial population size (**Fig. S13A**) and richness (**Fig. S13B** and **S14A**) were not significantly different between germinating and non-germinating seeds. Comparison of the relative abundance (RA) of seed-borne 16S ASVs in germinating and non-germinating seeds revealed significant increase (*P*< 0.01, log2FC >2) in RA of six and three ASVs in non-germinating seeds of bean and radish, respectively (**Fig. S13C**). However, according to the taxonomic affiliation at the genus level, none were found in common with the three and eight *gyrB* ASVs enriched in non-germinating seeds (**Fig. S14B**). Unfortunately, we were not able to recover representative bacterial strains for these enriched ASVs and therefore did not confirm the observed phenotype experimentally.

Once the seeds have germinated, the corresponding seedlings can be classified as normal or abnormal according to their morphologies (see Methods). By using this classification scheme, 22 normal and 22 abnormal bean seedlings were obtained as well as 40 normal and 10 abnormal radish seedlings (**Fig. 8A**). Initial bacterial population size and richness observed on seeds were not good predictors of seedling phenotypes (**Fig. 10A** and **10B**). Changes in RA of seed-borne ASVs were next investigated in bean seeds but not in radish as few seeds produced abnormal seedlings, which does not allow for a balanced distribution. Overall, the RA of six 16S ASVs and eight *gyrB* ASVs were significantly enriched (*P*<0.01, log2FC > 2) in seeds that produced abnormal seedlings (**Fig. 10C**). The taxonomic affiliation of four of these 16S and *gyrB* ASVs was in agreement at the genus-level (**Fig. 10C**). On the other hand, the RA of three 16S and six *gyrB* ASVs were significantly enriched in seeds that produced normal seedlings (**Fig. 10C**).

**Figure 10:**
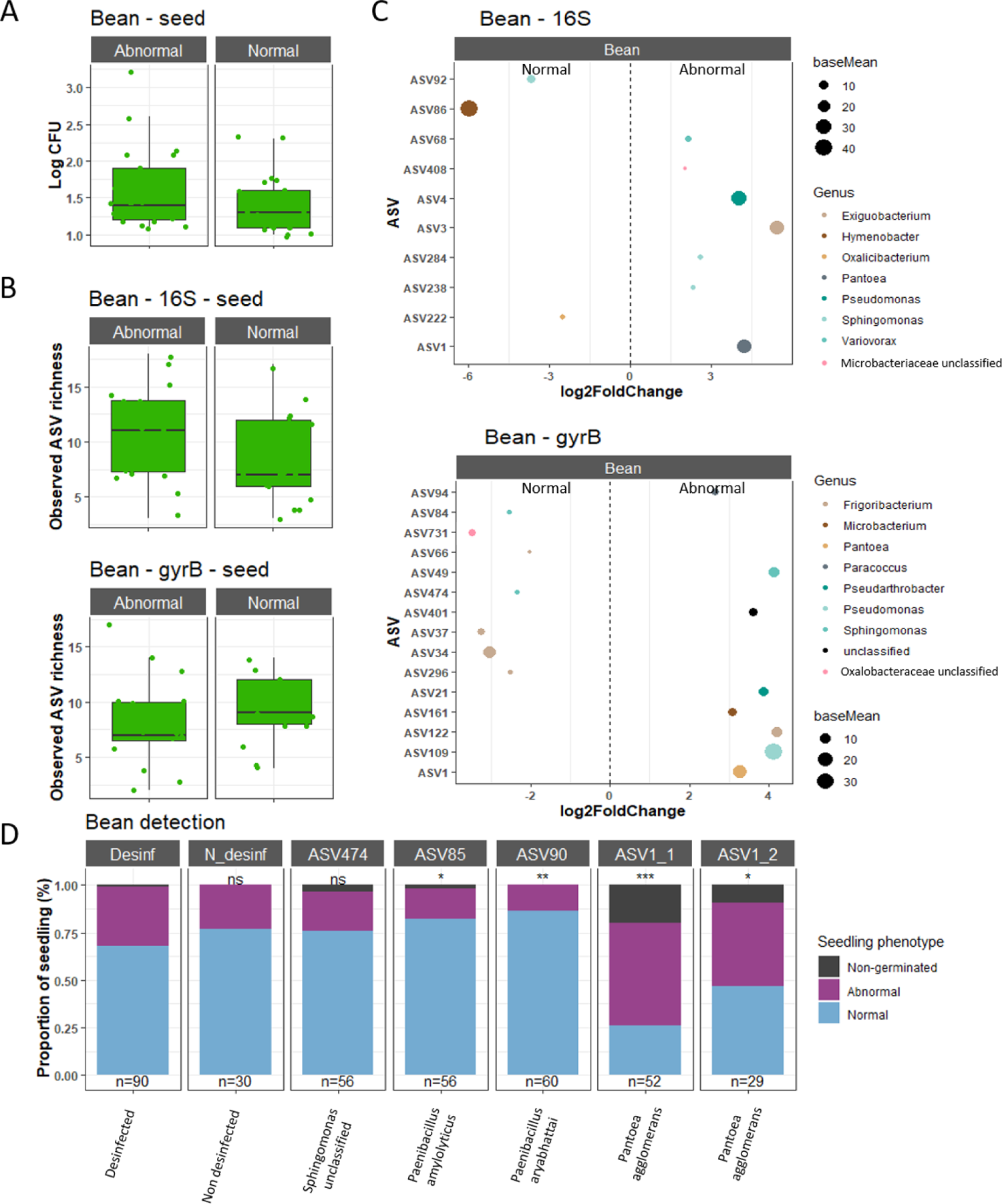
Impact of seed-borne ASVs on bean seedling phenotype. **A.** Correspondence between initial bacterial population size (log10 CFU) and seedling phenotype. **B.** Correspondence between observed richness on seeds and seedling phenotype. **C.** ASVs with significant changes in relative abundance (*P* < 0.01) in normal and abnormal seedlings. Colors corresponded to taxonomic affiliation at the genus level. **D.** Proportion of non-germinated seeds, normal or abnormal seedling after seed inoculation of bacterial strains. All seeds were primarily disinfected before inoculation. Strains were representative of ASVs significantly associated with normal (ASV85 and ASV90) and abnormal seedlings (ASV1) or associated with both phenotypes (ASV474). Statistical tests were performed with Chisquared test (ns = non-significant; * = *P* < 0.05; ** = *P* < 0.01; *** = *P* < 0.001).

To validate experimentally this phenotype, we isolated representative bacterial strains on the basis of their *gyrB* sequences and inoculated these strains on bean seeds (**Fig. 10D**). We selected strains representative of ASVs (i) significantly enriched in seeds that produced abnormal seedling (ASV1: *Pantoea agglomerans*), (ii) significantly enriched in seeds that produced normal seedling (ASV85: *Paenibacillus amylolyticus* and ASV90: *Bacillus aryabhattai*) and (iii) not significantly enriched (ASV474: *Sphingomonas* sp.). First, we validated that the seed surface-disinfection procedure employed did not impact seedling phenotype (**Fig. 10D**). As a control, seed inoculation of ASV474 did not induce any difference in seedling phenotype. In contrast seed-inoculation with two representative bacterial strains of ASV1 resulted in a significant (*P*<0.05) increase of abnormal seedling in comparison to surface sterilized seeds, mainly associated with rot symptoms. Inoculation of ASV85 and ASV90 increased the number of normal seedling (**Fig. 10D**). Hence, we confirmed the seedling phenotype expected based on the RA of ASVs.

## Discussion

The aim of this study was to assess the assembly of the seed microbiota during the seed development, its dynamics during seedling emergence and impact on seed vigor. Using a culture-enrichment procedure for performing community profiling at the individual seed level, we confirmed that seed transmission of bacteria is bottlenecked in bean and radish with the colonization of one dominant ASV of variable identity. Although abundance of seed-borne taxa were not predictive of efficient seedling transmission, their identities can significantly impact the bean seedling phenotype.

Community profile of a significant portion of collected seeds (bean 32%, radish 42%) could not be estimated as no CFU was detected on the culture medium employed, an observation already reported for seeds of several plant species^16^. The proportion of seed without detectable colony on TSA10 was negatively correlated with seed development stage in radish, while not impacted by seed development in bean. Independent of the development stage, spatial localization of seed was not a good estimator of CFU detection since seeds with or without detected colonies were often co-localized in the same fruit. Could the seeds without detectable colonies be considered as steriles? This is unlikely as CFUs were systematically obtained on each corresponding seedling, which justifies the use of seedlings for the detection of specific plant pathogens in the seed industry (a procedure called grow-out test^17^). The absence of CFU detection in some seeds could be explained by a localization in the internal tissues of the seed (i.e. on the surface of the embryo) which would not allow them to be obtained with the soaking procedures employed in this work. However, if this were the case, it is rather difficult to explain why these taxa would be located only in the embryo and not in the other cell layers of the seed since seed-borne bacteria are frequently visualized near the seed coat^24, 34^. Alternatively, it is likely that some seed-associated bacterial populations were in a state of viable but nonculturable (VBNC), which corresponds to a low metabolic activity state without formation of colonies on standard culture media. VBNC has been described in a number of seed-transmitted bacteria including *Clavibacter michiganensis* pv. *michiganensis*^32^ and *Acidovorax citrulli*^35, 36^. Why some populations enter the VBNC state in seeds and what are the signals that lift this state during germination-emergence are not yet known.

Overall, individual seeds were associated with a small number of taxa with a median of seven and four bacterial ASVs detected per individual seed of bean and radish, respectively. However, a dominant taxon representing 75 to 100% of all reads was associated with each individual seed, which could be considered as the primary symbiont hypothesis^18^. This notion of primary symbiont has been proposed for taxa associated with mature seeds at the end of the seed developmental process^18^. But did the early seed colonizers are systematically found on the mature seed? It seems that the answer to this question varies depending on the plant species considered. Although we acknowledge that the destructive sampling procedure employed in this work does not allow direct monitoring of the dynamics of the assembly of the microbiota during seed development, indirect evidence suggested a contrasting response for radish and bean. For radish, we did not observe significant changes in phylogenetic diversity of dominant taxa over time. Moreover, seed microbiota composition was also stable across the radish developmental stage, therefore suggesting that abundant taxa found during seed filling were mostly conserved during seed maturation. In contrast, increase in diversity during bean seed development together with a shift in taxonomic composition indicated that the early dominant taxa associated with bean seeds were progressively replaced during seed filling and maturation. This changes in community composition following colonization of a nearly sterile habitat (like seed) by micro-organisms is similar to primary succession^37^. Primary succession is influenced (*i*) the quantity of microorganism that can colonize a habitat and (*ii*) and changes in limiting resource through time^34^. Hence, the distinct microbial succession dynamics observed during bean and radish seed development is probably related to difference in dispersal and/or turnover of nutrient sources.

To estimate the relative importance of the different seed transmission pathways described to date (i.e. internal, floral and external^28^), atmosphere, flowers and stems were sampled throughout seed development. Overall, less than 20% of seed-borne taxa were detected in these three habitats, which probably reflected an undersampling of the source habitats in comparison to seeds. Although it is difficult to conclude on the relative importance of each habitat as a source of inoculum for the seed microbiota, we could see differences between the two plant species studied. Notably, the co-occurrence of ASVs in bean flower and seed was practically non-existent, which confirmed that seed transmission of bacteria through the floral pathway is weak in bean^7^. More interestingly, the variation in identity of seed-associated dominant ASV within the same sampling stage, especially between plants, appears to indicate that the local species pool varies. Variation in taxa composition is important for airborne communities and stem-associated communities, thus we hypothesize that both pathways might be important for bacterial transmission to the seed. However, airborne taxa have been demonstrated being a low source of microbial inoculum of leaves, fruits and flowers^38^. We therefore hypothesized that xylem sap could be the most important seed-transmission pathway. If this assumption is correct, then the observed infra-individual variation in seed microbiota composition (i.e. between leaf stages) implies a spatio-temporal dynamics of taxa in the xylem sap^39^.

Regardless of the origin of seed-borne taxa, what are the ecological processes involved in the assembly of the microbiota at the individual seed level? According to our neutral model, the relative importance of immigration rate was lower than local replacement rate for both plant species during seed development, even though for radish the relative contribution of immigration increased significantly during seed maturation. Local replacement could be either due to selection or ecological drift. For bean, selection is probably the main driver of community assembly during seed filling and maturation as the phylogenetic composition of seed communities were more similar than expected by chance. It is likely that this selection is due to host-filtering, notably through the production of chemical compounds and physical protective structures that developed during seed filling. From the primary succession dynamics observed during seed filling, the nature of the habitat is likely to change rapidly between seed filling and seed maturation, which appear to counterselect the Enterobacteriales. For radish, the relative importance of the different ecological processes analysed (selection, ecological drift and dispersal) is less clear. Indeed, βNTI values were scattered on both sides of the threshold −2 that is considered as significantly different from the null expectation. It is therefore likely that selection is less important in radish than in bean during seed development and that ecological drift is involved in the replacement rate observed. If this assumption is correct, this would be consistent with the results obtained by Rezki and coworkers, where drift was reported as the key ecological process during bacterial community assembly of mature seeds of radish.

The presence of taxa within mature seeds does not necessarily guarantee their transmission to seedlings^22^. In the absence of competition with other sources of inoculum such as the soil, we wondered whether taxa abundance on mature seeds is a strong predictor of seedling transmission? That is clearly not the case, since few seed dominant bacterial ASVs were dominant on seedling. This is in agreement with recent results obtained on rapeseed, where rare seed-borne or soil-borne taxa were efficiently transmitted on seedling^25^. As the initial inoculum size does not guarantee successful transmission to seedlings, then the difference in fitness between taxa (*Selection*) appears as a major driver of assembly of the seedling microbiota. In addition to host-filtering, competition for resources and spaces between taxa could be responsible for the selection of taxa in seedlings. Indeed, copiotrophic taxa were previously reported as dominant in seedling of bean and radish^24^. However, according to the taxonomic affiliation of ASVs, it seems that seedling-associated bacteria were not exclusively species with high growth rates (e.g. *Rhizobium* sp.), hence other traits are responsible for their selection.

Is seed vigour affected by the identity of the dominant taxon in seeds? In our study, overall seed microbiota structure and bacterial biomass did not influence seed vigour. However, we identified changes in RA of some specific ASVs in seeds that are correlated to non-germinating seeds or abnormal seedling. Moreover, we were able to validate the impact of seed-borne bacterial strains representative of these ASVs on seedling phenotype. Indeed seed-inoculation of two bacterial strains affiliated to *Paenibacillus amylolyticus* and *Bacillus* resulted in an increase of normal seedling Firmicutes are well known for their beneficial impact on plant growth. For instance seed-borne *Bacillus* species^40, 41^, and *Paenibacillus* species^42^ are known to significantly impact the seedling phenotype. In contrast, representative strains of *Pantoea agglomerans* (ASV1) caused an abnormal seedling phenotype, notably through root rot. *P. agglomerans* is a core seed microbiome taxon, highly abundant and observed in almost all plant species studied to date^43^. In bean, the occurrence of this taxa gradually decreased during seed filling and maturation. Therefore, symptoms caused on seedlings by *P. agglomerans* could occur when succession during seed development is not able to induce a low population level of this bacterial species.

## Conclusion

Individual seeds are associated with one dominant taxa of variable identity. These changes in identity are partly due to spatial heterogeneity not only between plants but also within the same plant. In addition, primary succession occurring during seed development are also responsible for these variations. Predicting these spatio-temporal changes at the seed level required an in-depth understanding of the processes involved in assembly of the seed microbiota. We showed that Selection is the main driver of community assembly but the nature(s) of this selective process remained to be explored. Research on microbial succession during the assembly of the seed microbiota and its seedling transmission will assist the design of seed microbiota engineering strategies through selection (i) of the most permissive seed transmission pathways and (ii) plant beneficial bacteria that enhance crop establishment.

## Methods

### Experiment 1 - Culture-based enrichment of seed-borne bacteria

Common bean (*Phaseolus vulgaris* L. var. Flavert) and radish (*Raphanus sativus* var. Flamboyant5) were at a density of eight bean plants and five radish plants per square meters at the IRHS experimental station (47°28’45.2’’N - 0°36’35.6’’W). Seeds were aseptically removed from fruits with sterile scalpel and tweezers under a laminar flow hood. Seed samples (*n*=24) were soaked in 2 ml of phosphate-buffered saline (PBS, Sigma-Aldrich) supplemented with Tween® 20 (0.05 % v/v, Sigma-Aldrich) per gram of fresh material. Seed soaking was performed at 4°C under constant agitation (150 rpm) for 2h30 and 16h for radish and bean, respectively. Half of the suspension was stored at −20°C, while the other half was serial-diluted and plated on 1/10 strength Tryptic Soy Agar (TSA10: 17 g.l^-1^ tryptone, 3 g.l^-1^ soybean peptone, 2.5 g.l^-1^ glucose, 5 g.l^-1^ NaCl, 5 g.l^-1^ K_2_HPO_4_, and 15 g.l^-1^ agar) supplemented with cycloheximide (50 µg.ml^-1^, Sigma-Aldrich). Plates were incubated at 18°C during 5 days. Bacterial mats were collected by adding 2mL of PBS Tween and stored at −20°C. DNA extraction was performed on (i) the frozen suspension and (ii) the collected CFU suspension with the NucleoSpin® 96 Food kit (Macherey-Nagel) following the supplier’s recommendations.

### Experiment 2 – Community assembly at the individual seed level

A second field trial was performed at the experimental station of the National Federation of Seed Multipliers (FNAMS, 47°28’012.42’’N - 0°23’44.30’’W, Brain-sur-l’Authion, France). Flowers from 5 plants at the blooming stage were collected, all flowers from one plant corresponding to one independent sample. Flowers were soaked in 2 ml of PBS Tween per gram of fresh material, then crushed in a lab blender (Stomacher, Mixwel, Alliance Bio Expertise) for 2 min. Individual seeds from five to seven plants of bean and radish were collected at different stages of the seed development (**Fig. 1**). A total of 96 and 144 seeds were sampled per radish and bean plant, respectively. Seed water content was estimated at each sampling stage on a bulk of 15 seeds by weighing this sample before and after drying (96°C for 3 days). Vascular flow of stem of each individual plant was collected as follows: a 2 cm section of stem from the bottom of the plant was cut and outer-surface-disinfected with 70° ethanol, then first layers of epidermis were removed using a sterile scalpel. The resulting piece of stem was incubated horizontally for 2.5 h in 2 mL of PBS Tween under constant agitation (200 rpm) at 4°C. Finally, airborne bacteria were collected at each sampling stage. To achieve this, a passive sampling scheme was performed with TSA10 plates placed at 10-20cm from the flowers. Six TSA10 plates were positioned homogeneously within the plot during 3 hours and then incubated at 18°C during 5 days.

Similarly, to experiment 1, all suspensions were serial-diluted and plated on TSA10. After 5 days of incubation at 18°C, the number of bacterial cells was estimated by counting the number of colony-forming units (CFU). We then determined a limit of quantification of 20 CFUs and an upper limit of quantification of 250 CFUs per plate^44^.DNA extractions were performed as described in experiment 1.

### Experiment 3 - Community dynamics during emergence

At the last sampling point of experiment 2 (D50 bean; D65 radish), additional mature seeds were collected in 2019 at the FNAMS experimental station. Thirty-two seeds per individual plants (*n*=5) were collected from eight fruits located at four different leaf heights. Four seeds were aseptically collected per fruit (**Fig. S2**). Seeds were soaked in PBS Tween. After collecting the resulting suspension, seeds were placed in sterile tubes filled with cotton moistened with 4mL of sterile water. Tubes were incubated in a growth chamber (photoperiod: 16h/8h, temperature 25-22°C) to obtain 4-day-old radish and 6-day-old bean seedlings. Seedlings were placed individually in a plastic bag, crushed and resuspended with 2 ml of sterile water. Suspensions of seeds and seedlings were serial-diluted and plated on TSA10 and DNA was extracted as previously reported. The sterility of our experimental system was assessed by collecting cotton from seedless tubes (n=30). No CFU was detected on TSA10 after 5 days of incubation at 18°C.

### Construction and sequencing of amplicon libraries

A first PCR amplification was performed with the primer sets 515f/806r^45^ and gyrB_aF64/gyrB_aR553^23^, which target the V4 region of 16S rRNA gene and a portion of *gyrB*, respectively. PCR reactions were performed with a high-fidelity Taq DNA polymerase (AccuPrimeTM Taq DNA polymerase Polymerase systemSystem, Invitrogen) using 5µL of 10X Buffer, 1µL of forward and reverse primers (gyrB_aF64/gyrB_aR553 [100µM]; 515f/806r [10µM]), 0.2µL of Taq and 5 µl of DNA. Cycling conditions for 515f/806r was composed of an initial denaturation step at 94°C for 3 min, followed by 35 cycles of amplification at 94°C (30 s), 50°C (45 s) and 68°C (90 s), and a final elongation at 68°C for 10 min. The cycling conditions for gyrB_aF64/gyrB_aR553 were identical with the exception of the annealing temperature (55°C instead of 50°C). Amplicons were purified with magnetic beads (Sera-MagTM, Merck). A second PCR amplification was performed to incorporate Illumina adapters and barcodes. PCR cycling conditions were as follows: denaturation at 94°C (2 min), 12 cycles at 94°C (1 min), 55°C (1 min) and 68°C (1 min), and a final elongation at 68°C for 10 min. Amplicons were purified with magnetic beads and pooled. Concentration of the pool was monitored with quantitative PCR (KAPA Library Quantification Kit, Roche). Amplicon libraries were mixed with 5% PhiX and sequenced with four MiSeq reagent kits v2 500 cycles (Illumina). A blank extraction kit control, a PCR-negative control and PCR-positive control (*Lactococcus piscium*, a fish pathogen that is not plant-associated) were included in each PCR plate.

### Microbial community analysis

All scripts and data sets employed in this work are available on GitHub (https://github.com/martialbriand/IRHS_EmerSys). Briefly, primer sequences were removed with cutadapt version 1.8^46^. Fastq files were processed with DADA2 version 1.6.0^47^. Chimeric sequences were removed with the removeBimeraDenovo function. Taxonomic affiliation of amplicon sequence variants (ASVs) was performed with a naive Bayesian classifier^48^ against the Silva 132 taxonomic training data or an in-house *gyrB* database available upon request.

Diversity analyses were conducted with Phyloseq version 1.32.0^49^. Sequences derived from 16S rRNA gene that were unclassified at the phylum-level, affiliated to Archaea, Chloroplasts and Mitochondria were removed. Since the primer set *gyrB*_aF64/*gyrB*_aR553 primers can sometimes co-amplified *parE*, a paralog of *gyrB*, the *gyrB* taxonomic training data also contained *parE* sequences. ASVs affiliated to *parE* or unclassified at the phylum-level were removed. Sequences were aligned with DECIPHER version 2.16.1 (Wright, 2016) and neighbor joining phylogenetic trees were constructed with Phangorn version 2.5.5^50^. Identification of sequence contaminants was assessed with decontam version 1.8.0^51^.

Alpha diversity metrics (richness and Faith’s phylogenetic diversity) were measured after rarefaction at 4,000 reads per sample for all datasets and both markers (**Fig. S15**). Changes in bacterial phylogenetic composition were estimated with weighted UniFrac distance^52^ on log10+1 transformed values. Permutational multivariate analysis of variance was performed with the adonis2 function of vegan 2.5.6^53^. Rank abundance curves were performed BiodiversityR 2.12.3 ^54^. Number of ASVs shared between specific contrasts were visualized with UpSetR 1.4.0^55^. Annotations of phylogenetic trees were performed with iTol 6.1^56^.

The relative importance of niche- or neutral-based processes in community assembly was estimated with abundance-based β-null deviation measures. Pairwise comparison of seed communities aggregated at the individual plant level was performed with β-mean-nearest taxon distance ^57^. Deviation from the null distribution was assessed with the β-nearest taxon index (βNTI^30^). βNTI values < −2 or > 2 indicates that the observed phylogenetic turnover between a pair of communities is primarily driven by Selection^30^.

### Analysis of species-proportion distributions

When considering the set of all ASV observed on a plant, only a few of them are detected in each seed. This is probably caused by censoring induced by the measurement process, but it probably also comes from an important spatial heterogeneity. We therefore proposed a dedicated approach to provide a species-proportion analysis in seeds at the plant level for the most abundant species (> 0.1% of the total ASV abundance). We used the neutral model of Solé and co-workers^29^ at the seed level, where all ASVs are assumed to be totally equivalent and driven by immigration (probability 0<mu<1) and internal random replacement (probability 0<C<1). Abundances in each seed follow a beta-binomial (BB) distribution with parameters *N_c_* (the local ASV population size in the seed), α_c_, β_c_ where S is the typical species pool size available for immigration in the vicinity of each seed and | is the ratio of the *per capita* immigration rate over the *per capita* replacement rate 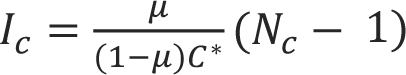 (Laroche et al. 2020). To obtain a distribution of seed-borne proportions at the plant level, we assumed that N followed a uniform distribution whose range [Nmin, Nmax] was determined as the 0.1-0.9 quantiles of the distribution of total ASV number in seeds at a given date. The other parameters 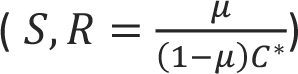 were assumed to be plant dependent. We computed from the data the proportions of ASVs in each seed at a given date on a given plant, evaluated the first three empirical moments of the mean at the plant level and performed a generalized moment estimation to obtain an estimated value for S, R and the expected I, as well as the value of the error criterion (see supplementary method for more details). All computations were coded in R, Wilcoxon test (resp. pairwise Wilcoxon rank test with Benjamini-Hochberg correction) was used for simple (resp. multiple) comparisons.

### Seedling phenotype

Seedling phenotype was estimated *via* rules established by the International seed testing association ^58^ (Briefly, a seedling was considered abnormal if at least 50% of the cotyledons or leaves were necrotic or rotten, if the hypocotyl or epicotyl were deformed, or if the root system was absent, stunted or rotten.

Changes in ASVs relative abundance were assessed with DESeq2 v 1.28 ^59^. Representative bacterial strains of ASVs significantly enriched in abnormal or normal seedling were selected from an in-house bacterial culture collection derived from bean seeds. Bean seeds (*Phaseolus vulgaris* var. Flavert), were surface-sterilized after 1 min sonication (40 Hertz), soaking for 1 min in 96° ethanol, 5 min in 2.6% sodium hypochlorite, 30 sec in 96° ethanol and rinsed 3 times with sterile water. The seeds were dried on paper and placed in a bacterial inoculum from a fresh 24-hour culture calibrated at DO = 0.01 (approximately 10^7^ CFU/mL). The seeds were then dried on paper. In order to check the inoculation of the seeds, half of the dried seeds were individually macerated in a pre-filled plate of PBS Tween. The other half of the seeds were placed in sterile tubes with cotton and moistened with 4mL of water, at the level of one seed per tube. Seedlings phenotypes were evaluated after 6 days in a growth chamber (photoperiod: 16h/8h, temperature 25-22°C). We also analyzed seedling phenotypes of non-disinfected seeds to see the impact of disinfection on seedling phenotype. To confirm the transmission of the inoculated strains, for each phenotype, we gathered seedlings in three plastic bags, one for non-germinated seeds, one for normal seedling and the last one for abnormal phenotypes. Batches were crushed and re suspended with 2 ml of sterile water per individual. Suspensions of seedlings were serial-diluted and plated on TSA 10%. After 5 days, we performed *gyrB* PCR amplification of representative members of each plate with gyrB_aF64/gyrB_aR553 primers. PCR products were sent to GenoScreen for Sanger sequencing.

## Acknowledgments

This work was supported by the French National Research Agency [ANR-17-CE20-0009-01]. The authors wish to thank Emmanuelle Laurent and Vincent Odeau (FNAMS) for field crop management as well as Muriel Bahut (ANAN platform, SFR Quasav) for amplicon sequencing.

## Author contributions

G.C. and M.Ba. designed the study program. G.C., B.L., M.S. and M.Ba wrote the manuscript. G.C., M.S., A.P., C.M. and B.M. designed and did the experimental part of the study. G.C., M.B and M.Ba. wrote all computational pipelines with the exception of the “*analysis of species-proportion distributions*”. The neutral model of this latter section was designed and written by B.L. All authors contributed to the editing and final revision of the manuscript.

## Data availability

The datasets supporting the conclusions of this article are available in the European Nucleotide Archive under the accession number PRJEB45079

## Code availability

All R scripts employed in this work are available on GitHub (https://github.com/martialbriand/IRHS_EmerSys)

## Supplementary method

### Species-proportion distribution analysis

We performed a qualitative analysis of the species-proportion distributions for both plant species. We focused on species thought to be actually representative of within seed microbiota, so microbial species whose abundance was less than 1% of the total abundance detected in one seed were discarded.

For each day and each plant, we computed the *empirical moments* of the observed ASV proportions. When considering the set of all ASV observed on a plant, only a few of them are detected in each seed. This is probably caused by censoring induced by the measurement process, but it probably also comes from spatial heterogeneity. This is why we computed the ASV proportions (*f*) for each seed related to the ASVs detected on that seed, and discarded the null frequencies for unobserved ASVs. Defining I_plant_ and J_seed_ as the set of seeds observed at a given date on a given plant and the set of ASV *observed at least one time* in this seed on that plant at the same date respectively, we obtain for each plant the following expressions for the first three empirical moments

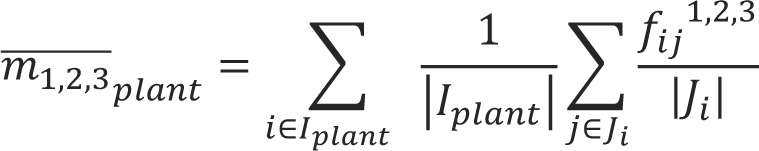

We compared these moments with those obtained with a neutral distribution derived from [Solé 2002]. These authors proposed the use of neutral distribution as a crude first order approximation of the species-abundance distributions in complex ecosystems. This distribution is the stationary distribution of a neutral model, involving local species communities (for us, a microbial community associated to a seed in a plant) and a “local” regional pool (for us, a plant associated community in the vicinity of a seed). Here, neutrality is taken in the sense of [Laroche et al 2020], that is (i) local per capita rates of birth and death are identical for all individuals, whatever their species, (ii) immigration in the local community occurs through random sampling of individuals from the regional pool, (iii) all these rates depend on the local community size only. Moreover, it is assumed in this model that the local community is saturated (constant size *N*>0), and the typical local species pool size (the number of ASV available for immigration from the plant and its close environment in the vicinity of a seed) is *S*>0.

Two phenomena are accounted for, immigration (probability |, such that 0 < | < 1) and local competition, modelled through a parameter *C*, representing the percentage of ASV with which an ASV competes, with equal probability to win. This parameter is also called the connectivity of the local community. The corresponding probability to win is *C*^*^ = 1 − (1 − *C*)^2^.

Standard results in ecological theory (see [Laroche et al. 2020] for a review) lead to the definition of the ratio of the per capita immigration rate (|) over the per capita replacement rate ((1 − |)*C*^∗^/(*N_c_* − 1)) in a community (seed) c of size *N_c_*.

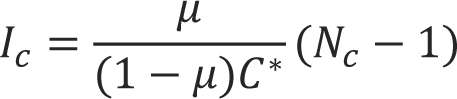

This is the relevant ecological parameter in these models, measuring the relative strength of immigration and internal competition or selection dynamics that shape the species abundance in the community, together with the local species pool size S. The species abundance distribution *P* is then given by a Beta-binomial distribution with parameters *N_c_*, α_c_, β_c_ expressed as

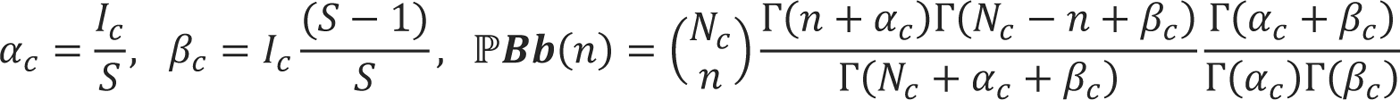

### We modified this model to compute the species-abundance distribution at the plant level for a given day

To this end, we hypothesized that for seeds in which at least one microbe was detected, the distribution of total number of detected ASVs in all seeds was an acceptable proxy of the distribution of *N_c_* accounting for natural variability and variability induced by the sampling process, but ignoring cultivation bias. Moreover, after inspection of the data, we assumed that for each observation date d

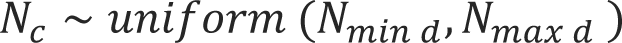

where *N*_min d_, *N*_max d_ depends on the sampling date but are the same for all plants.

Therefore, the marginal species abundance distribution at date d for seeds of plant p is given by

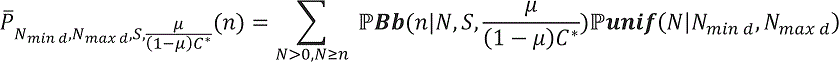

where ℙ**Bb** and ℙ**unif** are the Beta-binomial and uniform (on integer) probability functions.

We may notice that the model actually depends on the ratio 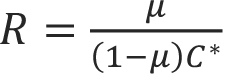 and the parameter *S* and *N*_min d_, *N*_max d_. As we actually only observe proportions of ASVs present within each seed, we compute the three first moments of positive proportion distribution at the plant level for this model:

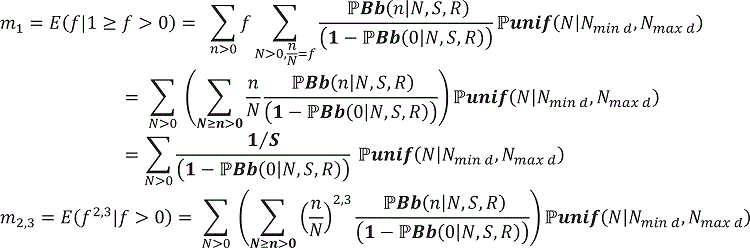

Based on this model, for each date we used the total number of ASVs to choose *N*_min d_, *N*_max d_ separately. After plotting the quantiles for each date, we observed that we could reasonably assume a uniform distribution on the interval corresponding to the 0.1 and 0.9 quantiles. Then, for a given set of parameters we simulated 1000 values of *N* and computed the moments according to the above formula.

We performed a **generalized moment estimation** of the parameters (*S, R*) to fit the observed moments for each plant. For this we screened the following parameter grid (whose range was selected by preliminary trial):

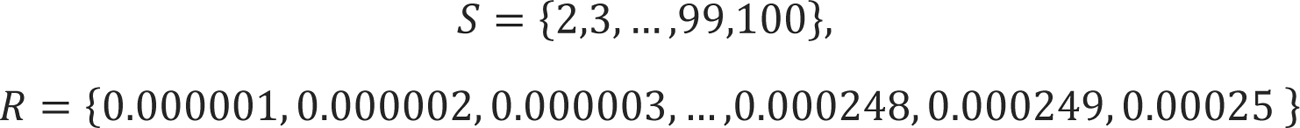

and selected for each plant the set of parameters that minimize the quantity

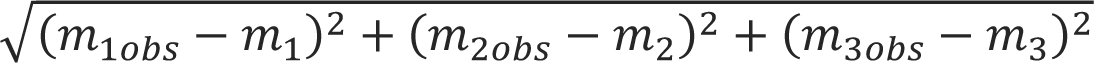

We also tested a weighted version of this criterion, under the form

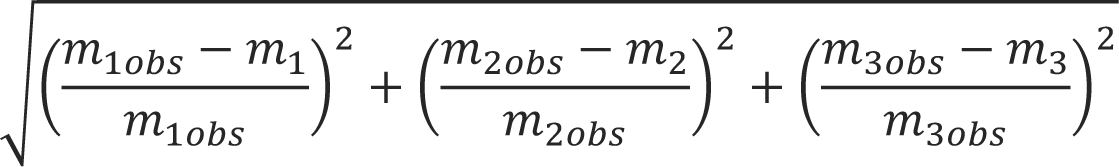

The empirical and estimated probability densities of proportions for each day and each plant were tested and plotted for bean and radish with 16S marker (**FigSs1**).

---

**Figure S1:**
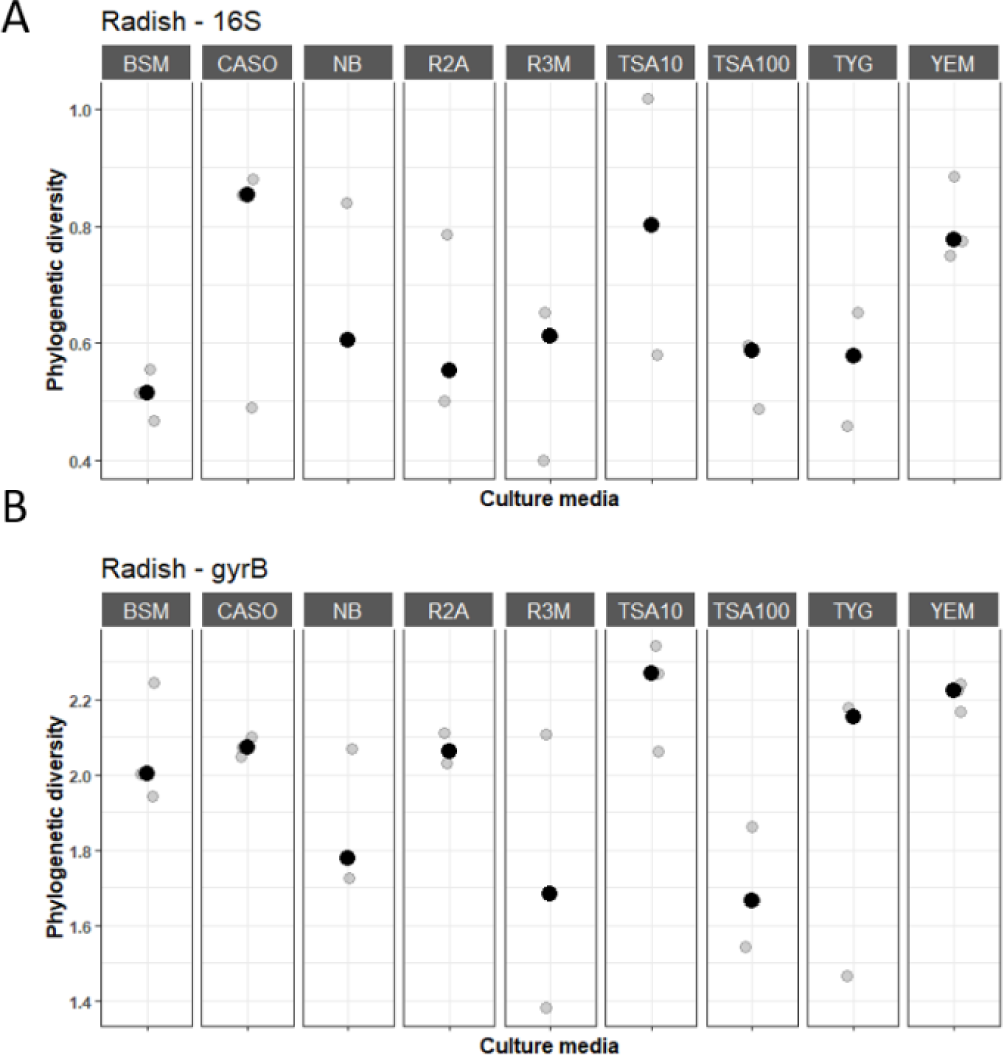
Phylogenetic diversity of seed bacterial communities after culture-enrichment on different synthetic media. Bacterial suspensions derived from radish seeds after soaking in PBS Tween (three subsamples of 1,000 seeds each) were diluted and plated on the following nine culture media: <u>b</u>acterial <u>s</u>creening <u>m</u>edium 523 (BSM); <u>ca</u>sein peptone <u>so</u>ybean flour peptone agar (CASO); <u>n</u>utrient <u>b</u>roth agar (NB); <u>R</u>easoner’s <u>2A</u> (R2A); <u>r3 m</u>edium (R3M) (Dione et al., 2016); tryptic <u>s</u>oy <u>a</u>gar 1/10 strength (TSA10); tryptic <u>s</u>oy <u>a</u>gar (TSA100); tryptone <u>y</u>east extract <u>g</u>lucose (TYG) and <u>y</u>east <u>e</u>xtract <u>m</u>annitol agar (YEM). Phylogenetic diversity (Faith’s PD) of bacterial communities was estimated with the v4 region of 16S rRNA gene (**A**) and *gyrB* (**B**). Black circles represented the average phylogenetic diversity of three replicates, which are displayed in grey.

**Figure S2:**
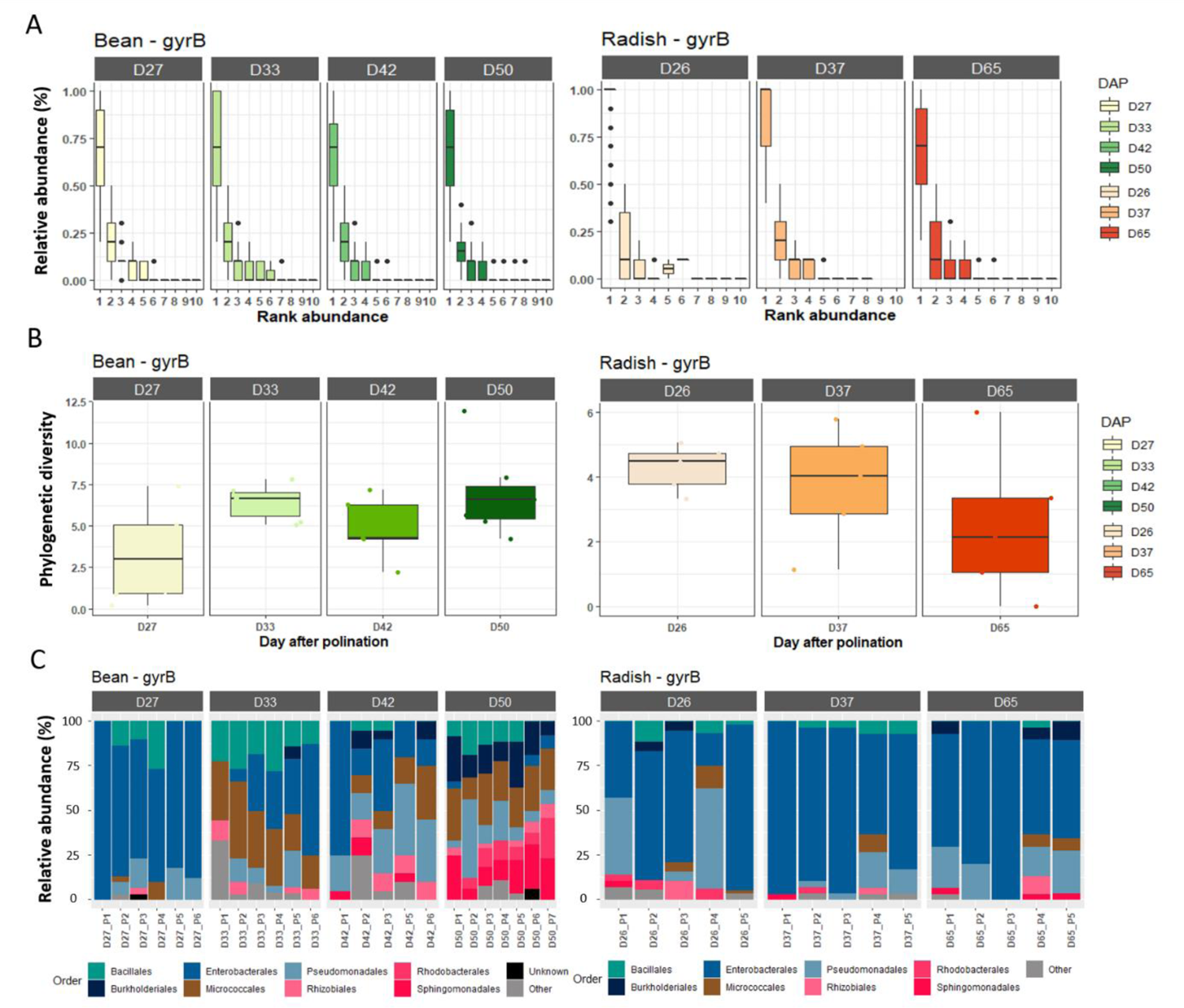
Dominant bacterial taxa within the seed microbiota based on *gyrB*. **A.** Rank abundance curves of bacterial ASVs associated with individual seeds. **B.** Faith’s phylogenetic diversity of rank1 ASV agglomerated at the plant level. **C.** Relative abundance of bacterial Orders calculated from rank 1 ASV agglomerated at the plant level. The names in each frame correspond to the stages sampled during seed development. The letters (a, b, c) indicated the significance level (*P* < 0.01).

**Figure S3:**
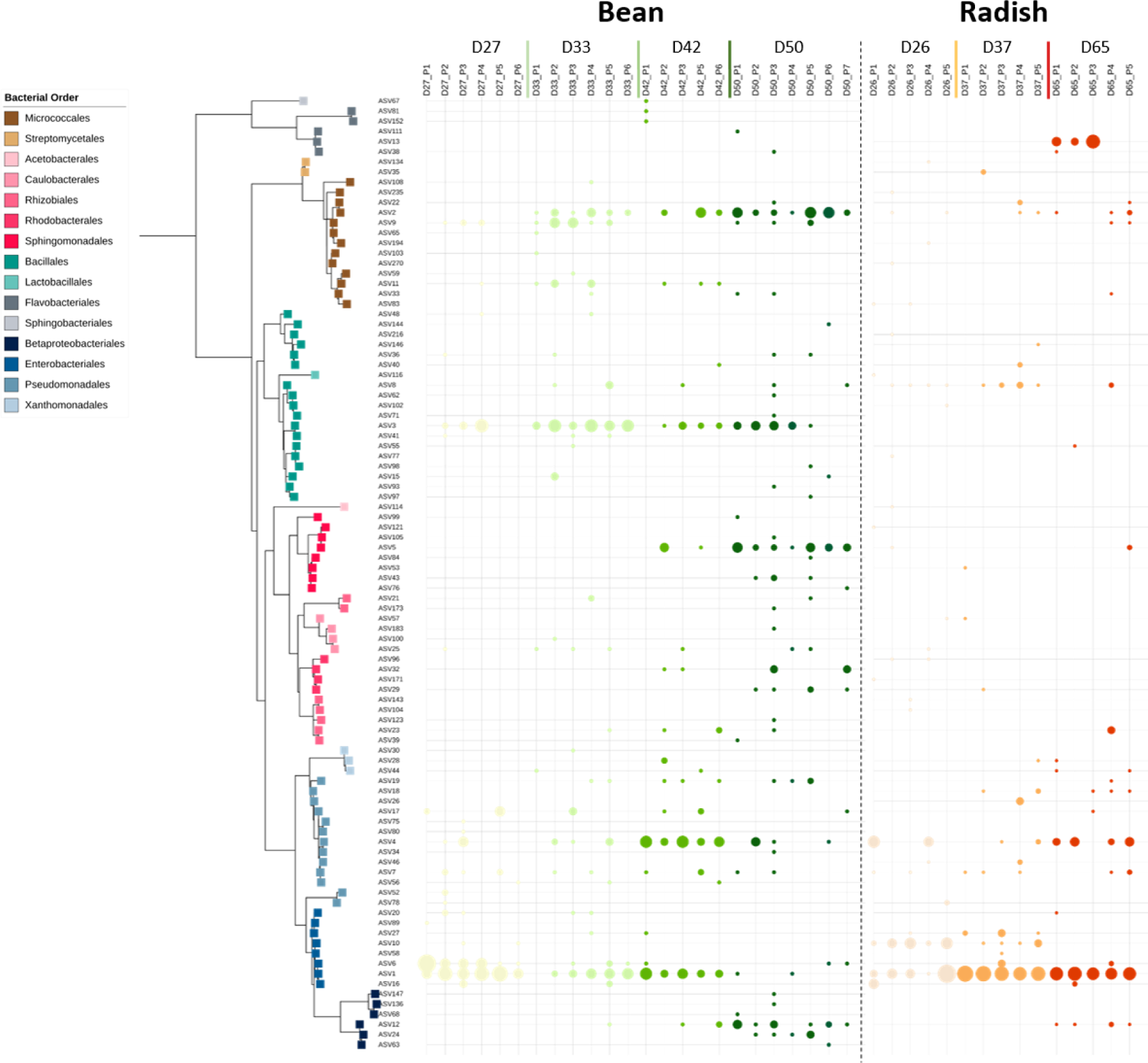
Phylogenetic diversity of seed-borne dominant taxa based on 16S. Neighbor-joining phylogenetic tree of dominant 16S ASV. Occurence of seed-borne ASVs within a specific plant is represented by a circle whose size is proportional to its relative frequency. Tips labels represented taxonomy at the Order level.

**Figure S4:**
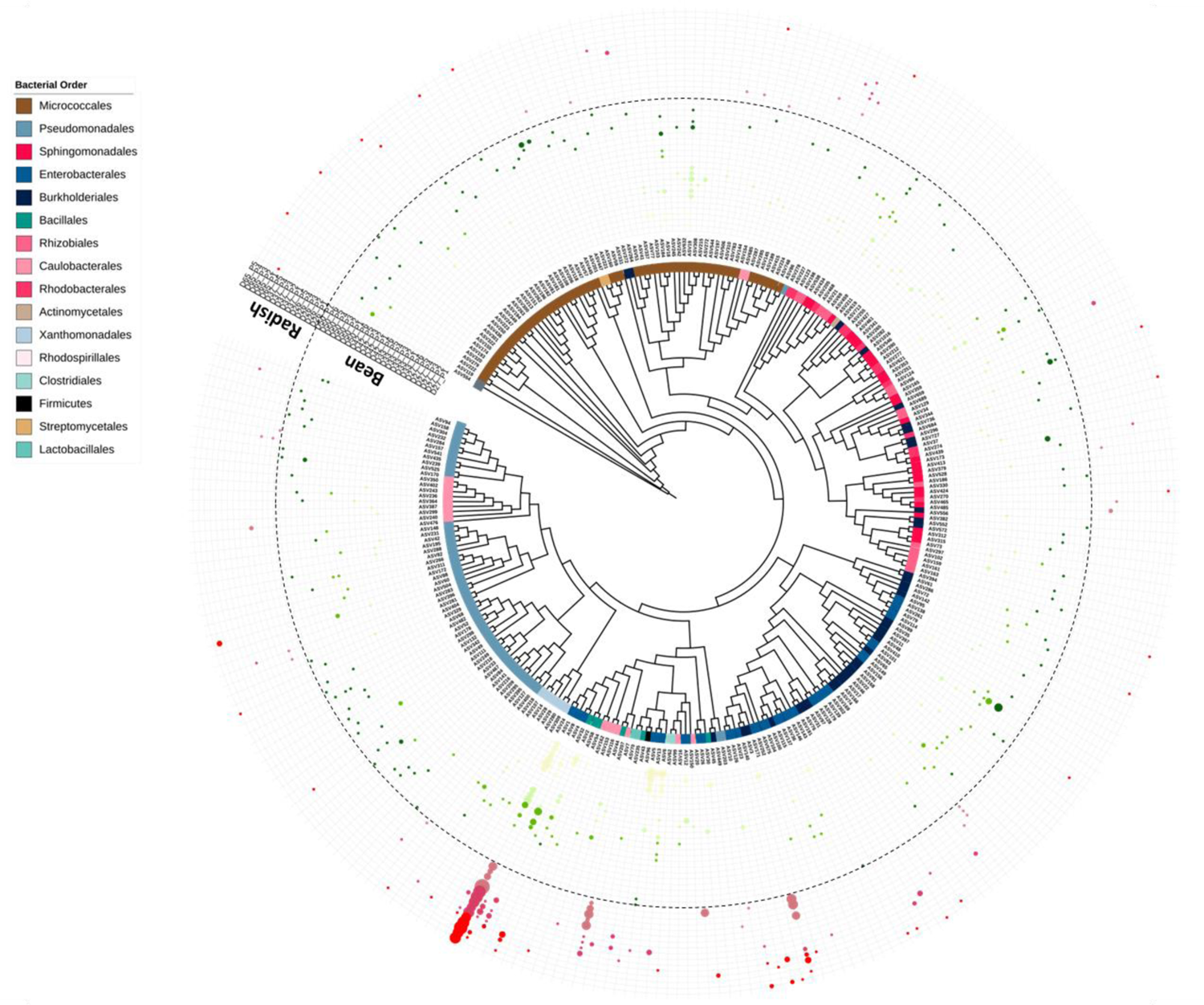
Phylogenetic diversity of seed-borne dominant taxa based on *gyrB*. Neighbor-joining phylogenetic tree of dominant *gyrB* ASV. Occurence of seed-borne ASVs within a specific plant is represented by a circle whose size is proportional to its relative frequency. Tips labels represented taxonomy at the Order level.

**Figure S5:**
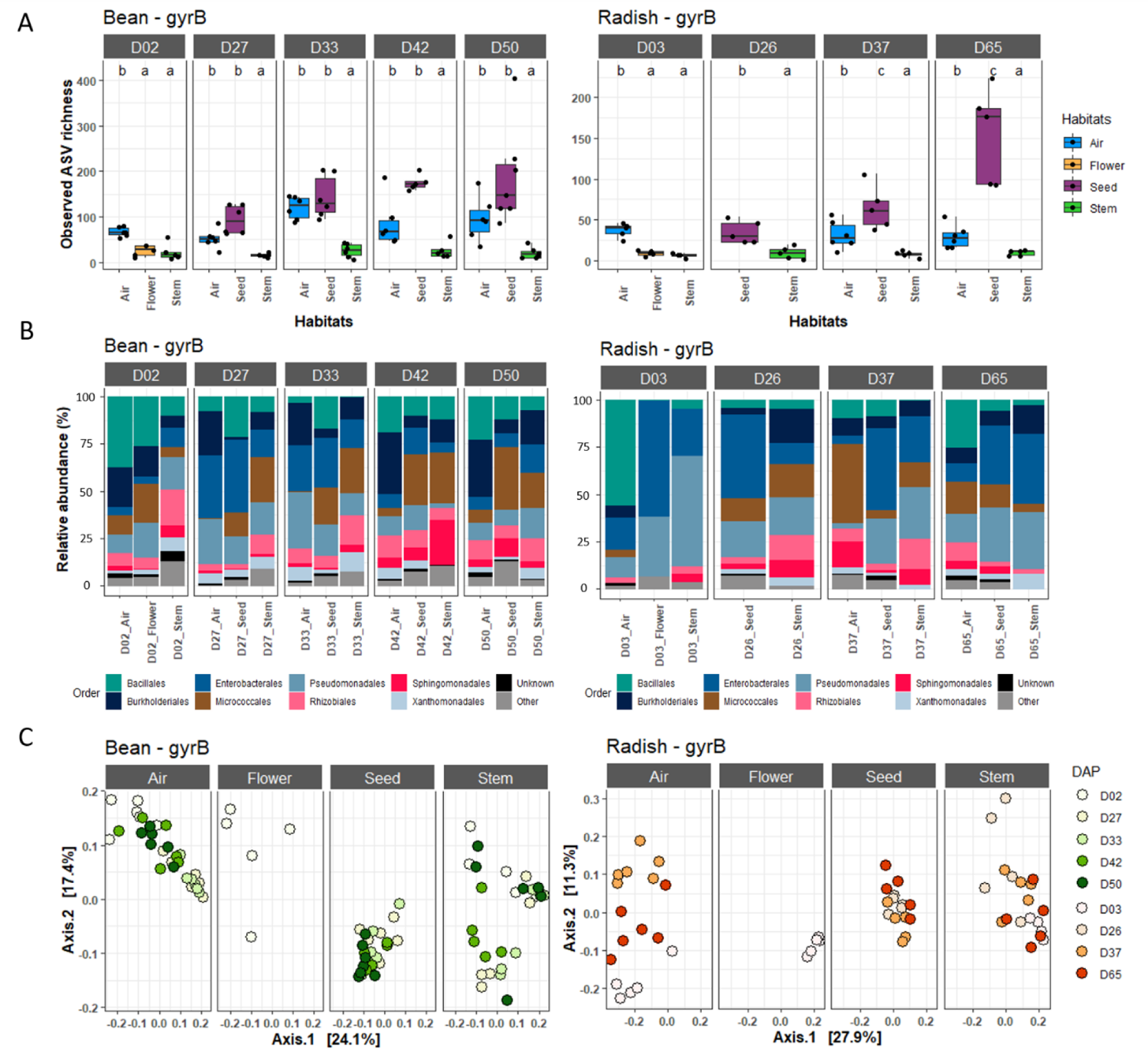
Structure of microbiota associated with flowers, stems and atmosphere. **A.** Observed richness for each habitat. The letters (a, b, c) indicated the significance level (*P* < 0.01). **B.** Relative abundance of bacterial Orders for each habitat. **C**. Variation in bacterial phylogenetic composition - as assessed with weighted UniFrac distances.

**Figure S6:**
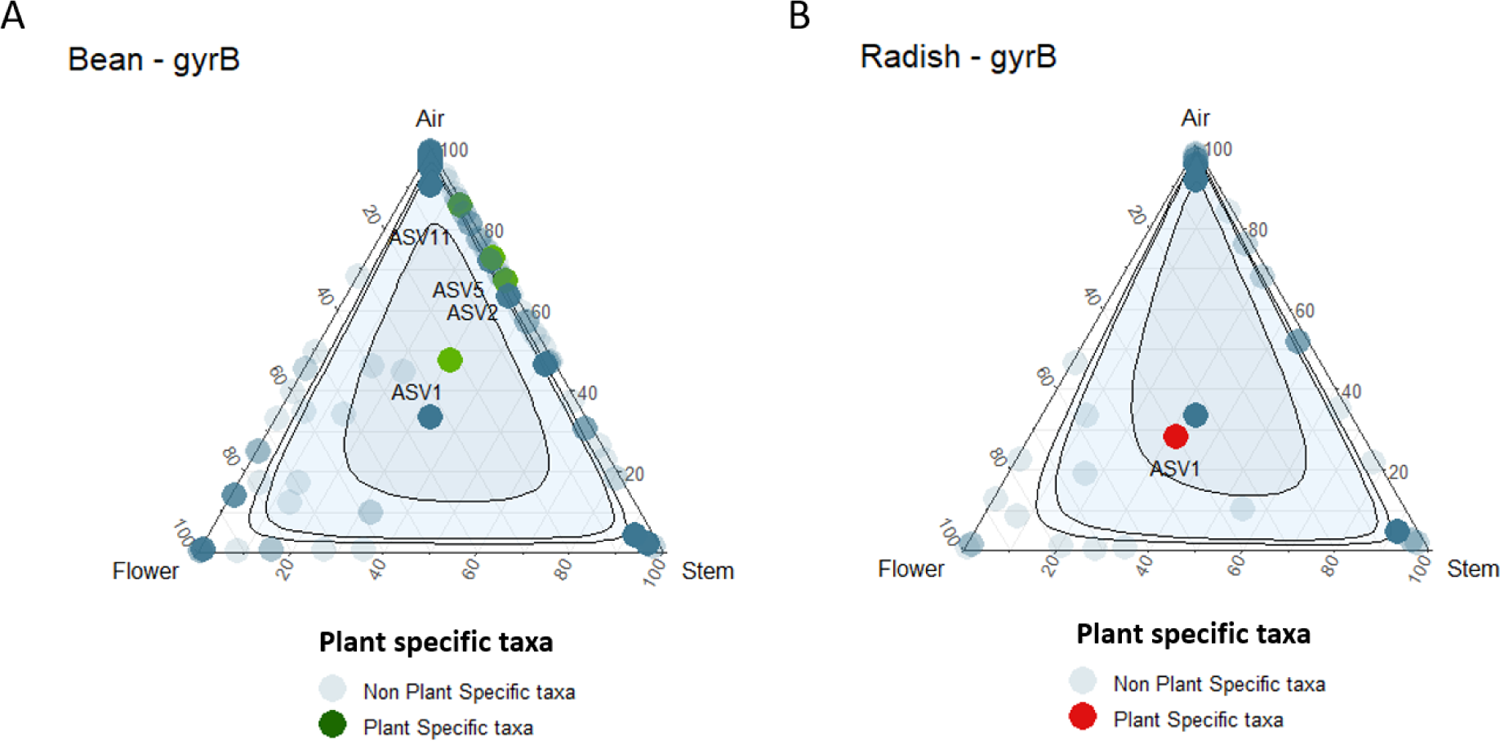
Origin of dominant seed-borne taxa. **A and B.** Detection of dominant seed-borne ASVs (*gyrB*) in other sampled habitats: flowers, stems and atmosphere. Intervals of confidence are indicated by a blue colour gradient ranging from 0.50 to 0.95. ASVs significantly associated with a specific plant (see **Table S1**) are highlighted in green (**A**) and red (**B**) for bean and radish. Of note, ASV20 is not represented for radish because it was not detected in any sampled habitats.

**Figure S7.**
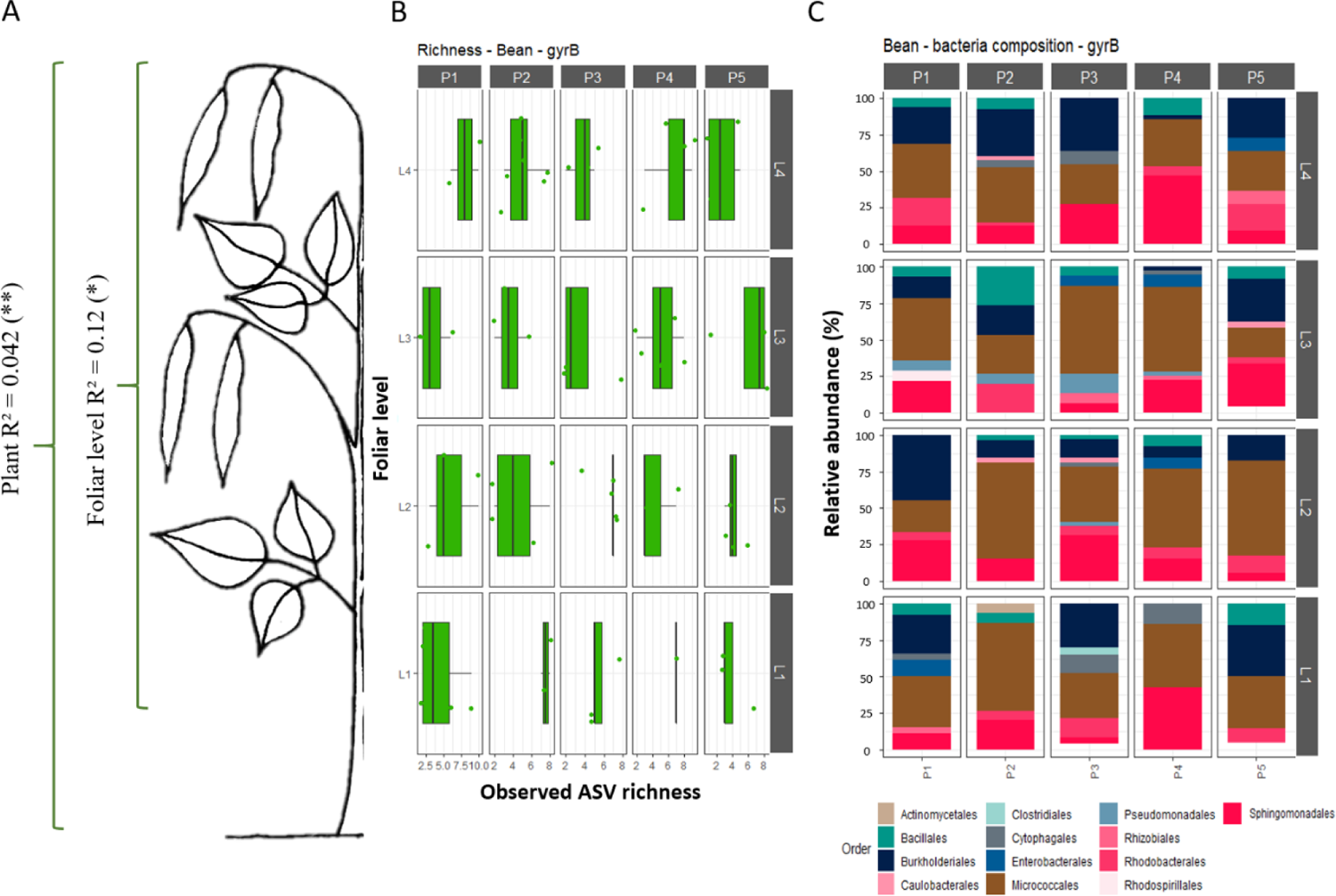
Influence of foliar level on the structure of bean seed microbiota. **A.** Variation in seed bacterial phylogenetic composition (weighted Unifrac distance) according to individual plants and foliar level inside each plant (PERMANOVA; * = *P* < 0.05; ** = *P* < 0.01). **B.** Average richness of seed microbiota per foliar level inside each plant. **C.** Relative abundance of bacterial Orders per foliar level. Foliar levels are represented as a gradient from the bottom of the plant (L1) to the top of the plant (L4), y-axis. All analyses were performed with *gyrB*.

**Figure S8:**
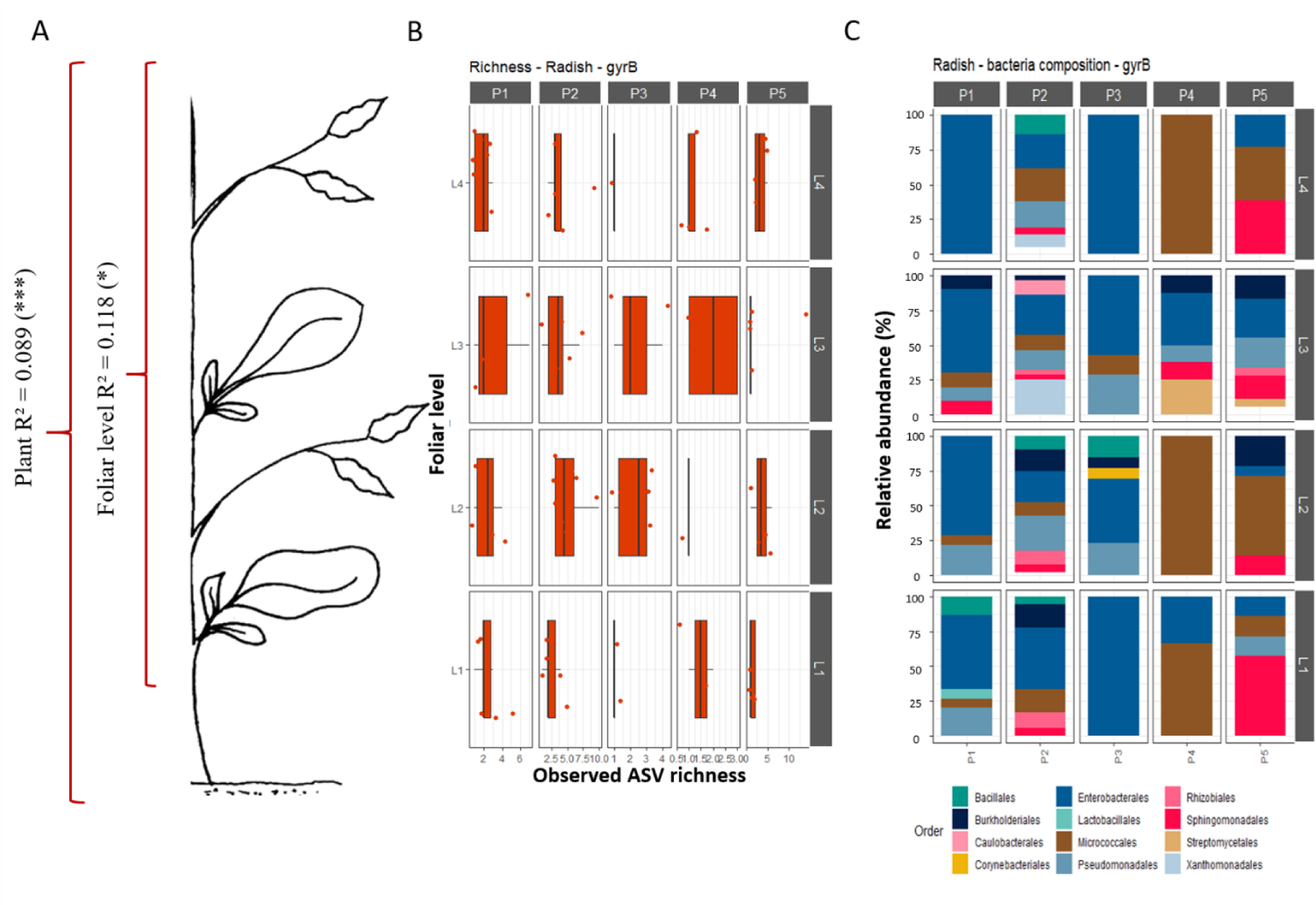
Influence of foliar level on the structure of radish seed microbiota. **A.** Variation in seed bacterial phylogenetic composition (weighted Unifrac distance) according to individual plants and foliar level inside each plant (PERMANOVA; * = *P* < 0.05; ** = *P* < 0.01). **B.** Average richness of seed microbiota per foliar level inside each plant. **C.** Relative abundance of bacterial Orders per foliar level. Foliar levels are represented as a gradient from the bottom of the plant (L1) to the top of the plant (L4), y-axis. All analyses were performed with *gyrB*.

**Figure S9:**
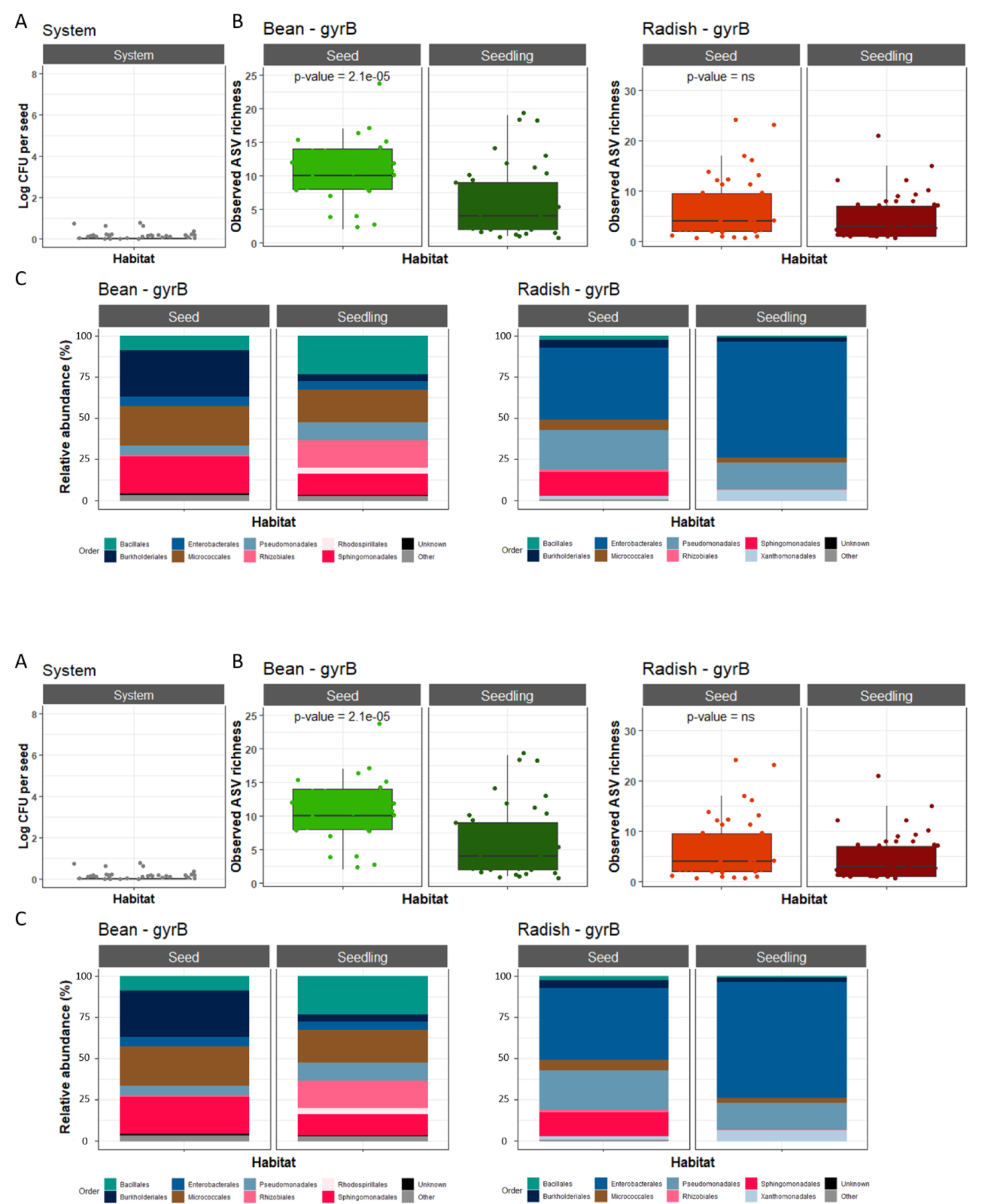
Comparison of seed and seedling microbiota profiles. **A.** Log10 CFU detected after seven days in our experimental system composed of tubes filled with cotton and sterile water. **B.** Average richness (estimated with *gyrB* ASVs) in seed and seedling. **C.** Taxonomic profile (Order level - *gyrB*) of seeds and seedlings microbiota. *P*-values were derived from a Wilcoxon rank sum test.

**Figure S10:**
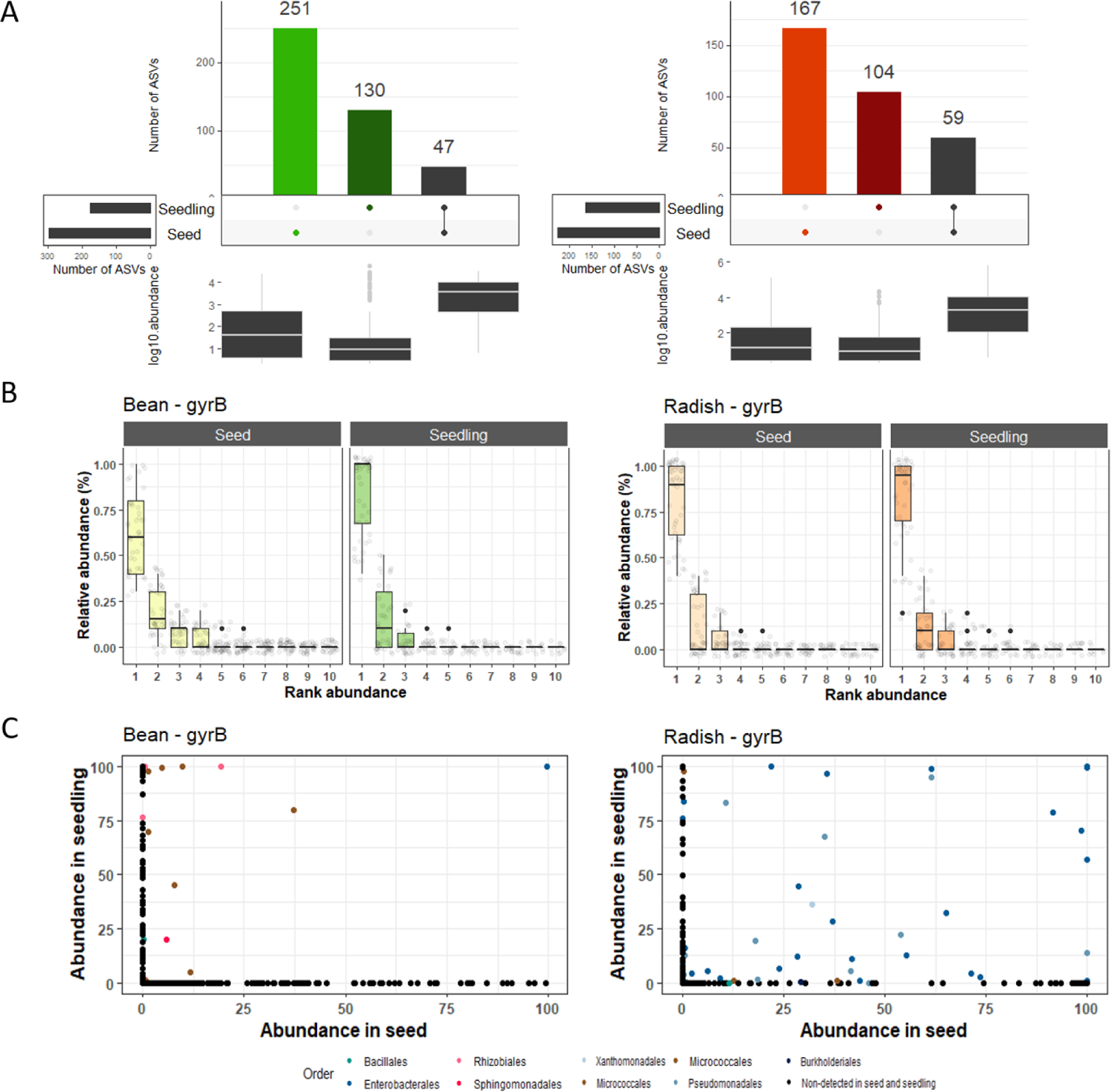
Seed to seedling transmission. **A.** Number of *gyrB* ASVs specifically detected on seeds (light color), seedlings (dark color) or both habitats (black). **B.** Rank abundance curves of *gyrB* ASVs in individual seeds and seedlings. **C.** Comparison of *gyrB* ASV relative abundance in seed and seedling. Each dot corresponded to one ASV, which is colored according to its taxonomic affiliation at the Order level.

**Figure S11:**
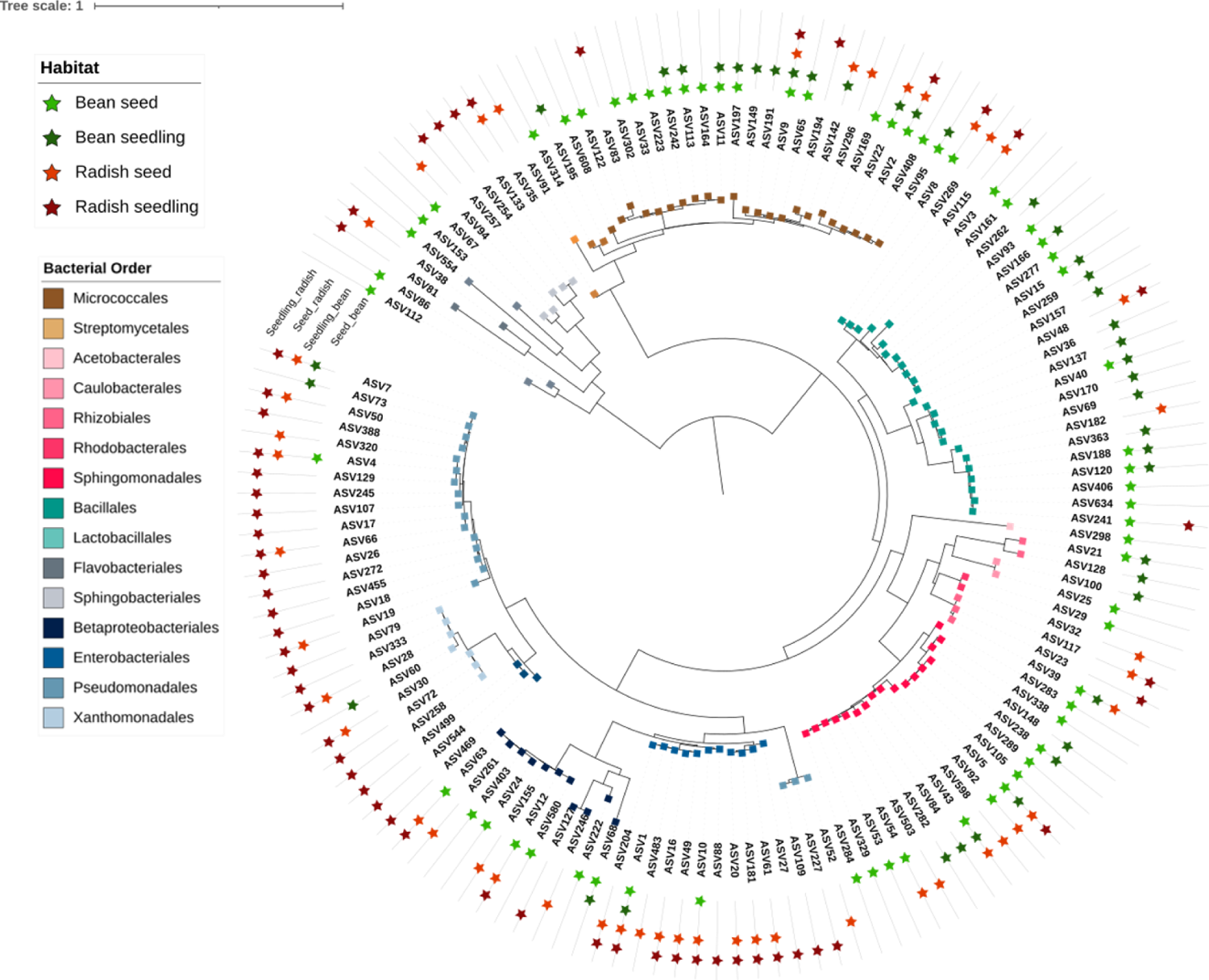
Phylogenetic diversity of 16S ASVs associated with seeds and seedlings. Neighbor-joining phylogenetic tree of bacterial ASVs. Stars represent ASV present in bean seed (light green), bean seedling (dark green), radish seed (light red) or radish seedling (dark red). Colors represent taxonomy at the Order level.

**Figure S12.**
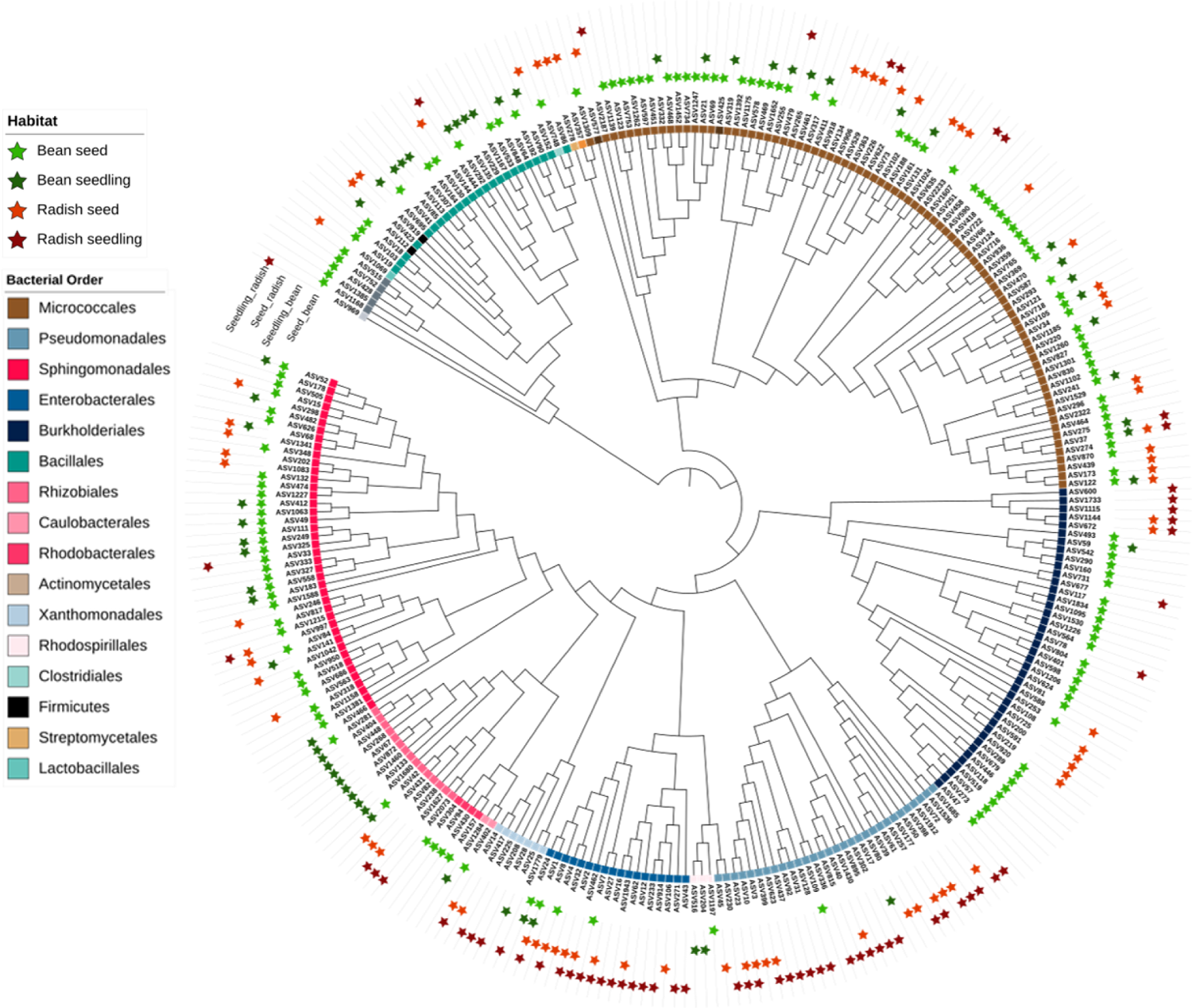
Phylogenetic diversity of *gyrB* ASVs associated with seeds and seedlings. Neighbor-joining phylogenetic tree of bacterial ASVs. Stars represent ASV present in bean seed (light green), bean seedling (dark green), radish seed (light red) or radish seedling (dark red). Colors represent taxonomy at the Order level.

**Figure S13.**
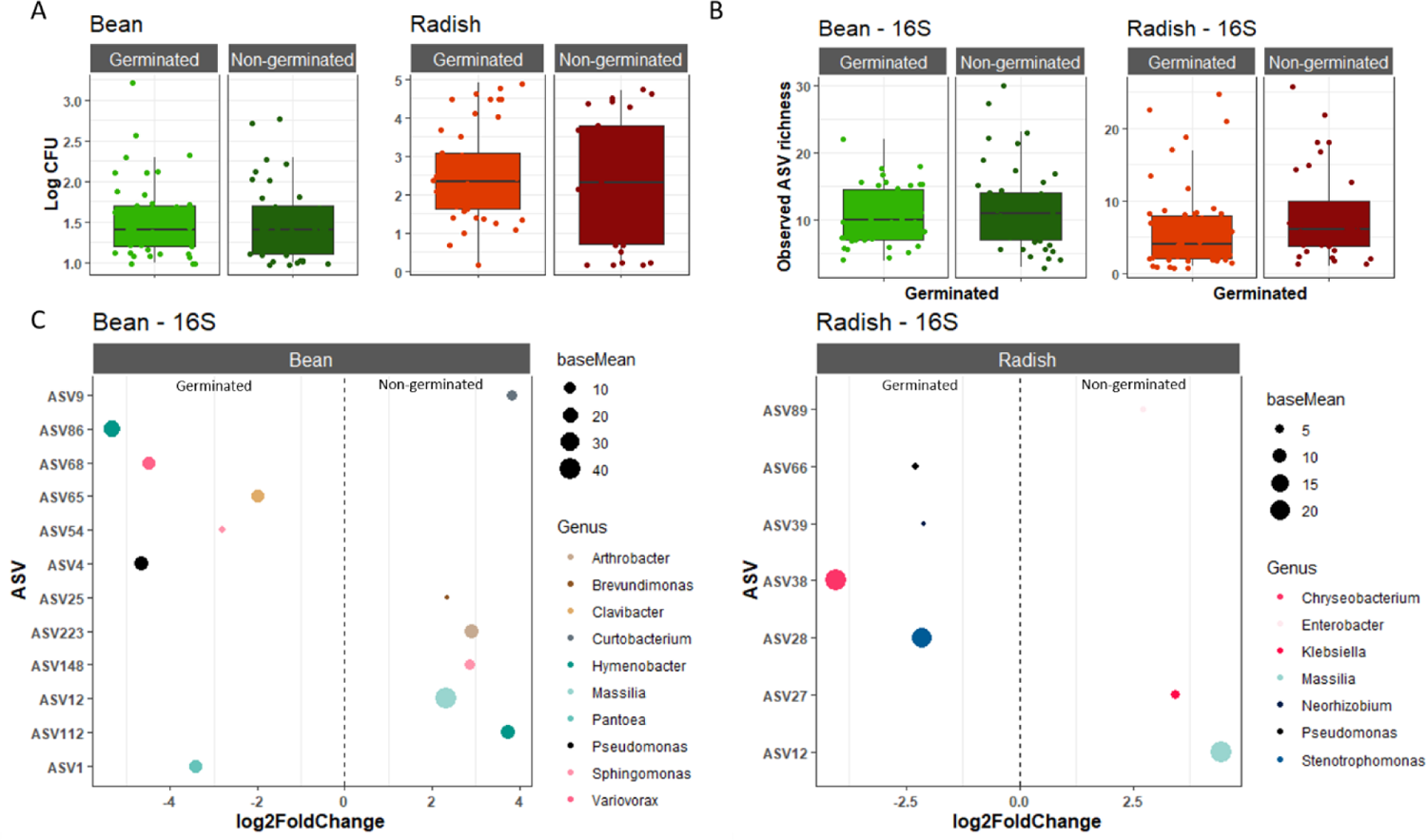
Differences in seed microbiota profile (16S) between germinated and non-germinated seeds. **A.** Correspondence between initial bacterial population size (log10 CFU) and seed germination. **B.** Average richness in germinated and non-germinated seeds. **C.** ASVs significantly (*P* < 0.01) enriched in germinated seeds (left) or non-germinated seeds (right). ASVs are colored according to their taxonomic affiliations at the genus level.

**Figure S14.**
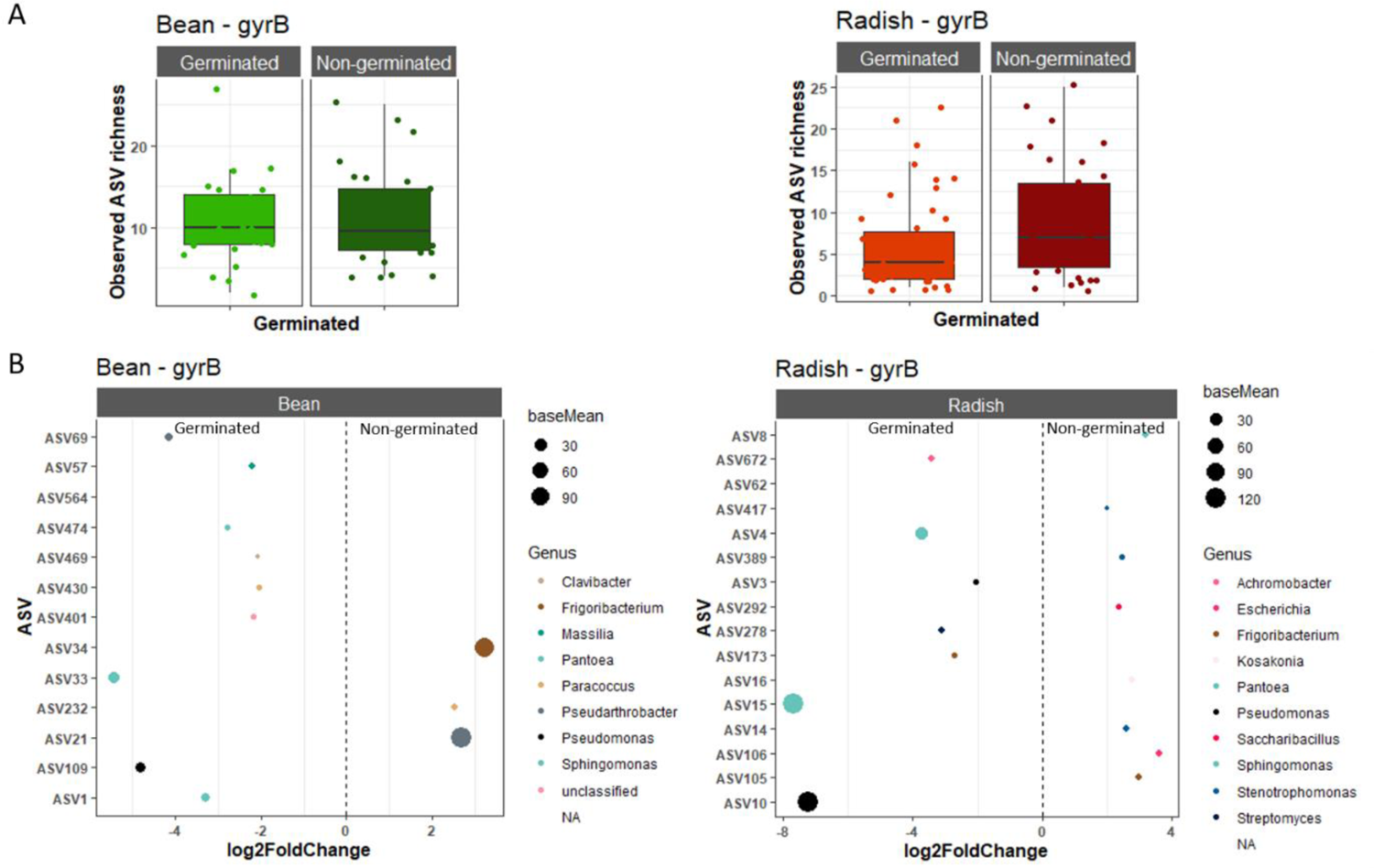
Differences in seed microbiota profile (*gyrB*) between germinated and non-germinated seeds. **A.** Average richness in germinated and non-germinated seeds. **B.** ASVs significantly (*P* < 0.01) enriched in germinated seeds (left) or non-germinated seeds (right). ASVs are colored according to their taxonomic affiliations at the genus level.

**Figure S15.**
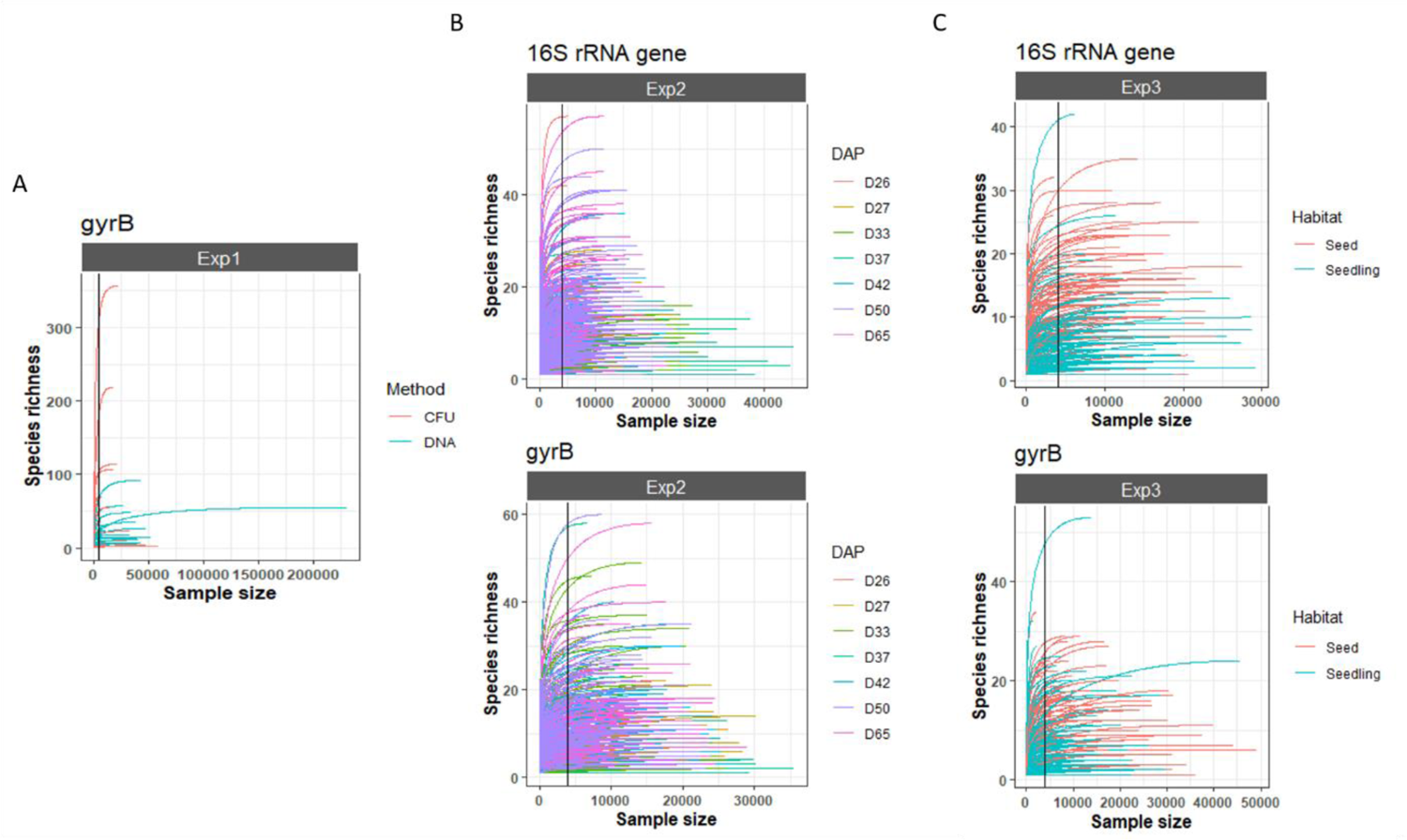
Rarefaction curves for all data sets. **A.** data set 1 **B.** data set 2 **C.** data set3. Black vertical lines correspond to a rarefaction threshold of 4,000 reads.

**Table S1:** ASVs significantly associated with one individual plant at each stage of development.

| Plant species | Plant | 16S rRNA gene |  |  |  |  | gyrB |  |  |  |  |
| --- | --- | --- | --- | --- | --- | --- | --- | --- | --- | --- | --- |
|  |  | ASV | Taxonomy | Frequency | Residual | Stem (RA%) | ASV | Taxonomy | Frequency | Residual | Stem (RA%) |
| Bean | D27_P1 | ASV6 | Erwiniaceae | 0,73 | 4,95 | 24,52 | ASV5 | Erwiniaceae | 0,47 | 3,27 | 58,23 |
| Bean | D27_P2 | - | - | - | - | - | ASV2 | *P. agglomerans* | 0,20 | 3,13 | nd |
| Bean | D27_P5 | - | - | - | - | - | ASV1 | *P. agglomerans* | 0,64 | 3,57 | 45,18 |
| Bean | D33_P6 | - | - | - | - | - | ASV1 | *P. agglomerans* | 0,44 | 3,82 | 20,09 |
| Bean | D50_P2 | ASV4 | *Pseudomonas* | 0,35 | 4,12 | nd | ASV11 | *P. putida* | 0,44 | 7,02 | nd |
| Radish | D26_P1 | ASV4 | *Pseudomonas* | 0,38 | 3,87 | 8,98 | - | - | - | - | - |
| Radish | D26_P1 | ASV16 | Erwiniaceae | 0,24 | 4,70 | nd | ASV20 | *P. ananatis* | 0,25 | 4,66 | nd |
| Radish | D26_P4 | ASV4 | *Pseudomonas* | 0,44 | 3,53 | 99,97 | - | - | - | - | - |
| Radish | D26_P5 | ASV1 | *Pantoea* | 0,71 | 3,67 | nd | ASV1 | *P. agglomerans* | 0,77 | 3,73 | nd |
| Radish | D65_P3 | ASV13 | *Chryseobacterium* | 0,53 | 4,70 | NA | - | - | - | - | - |

**Figure Ss1:**
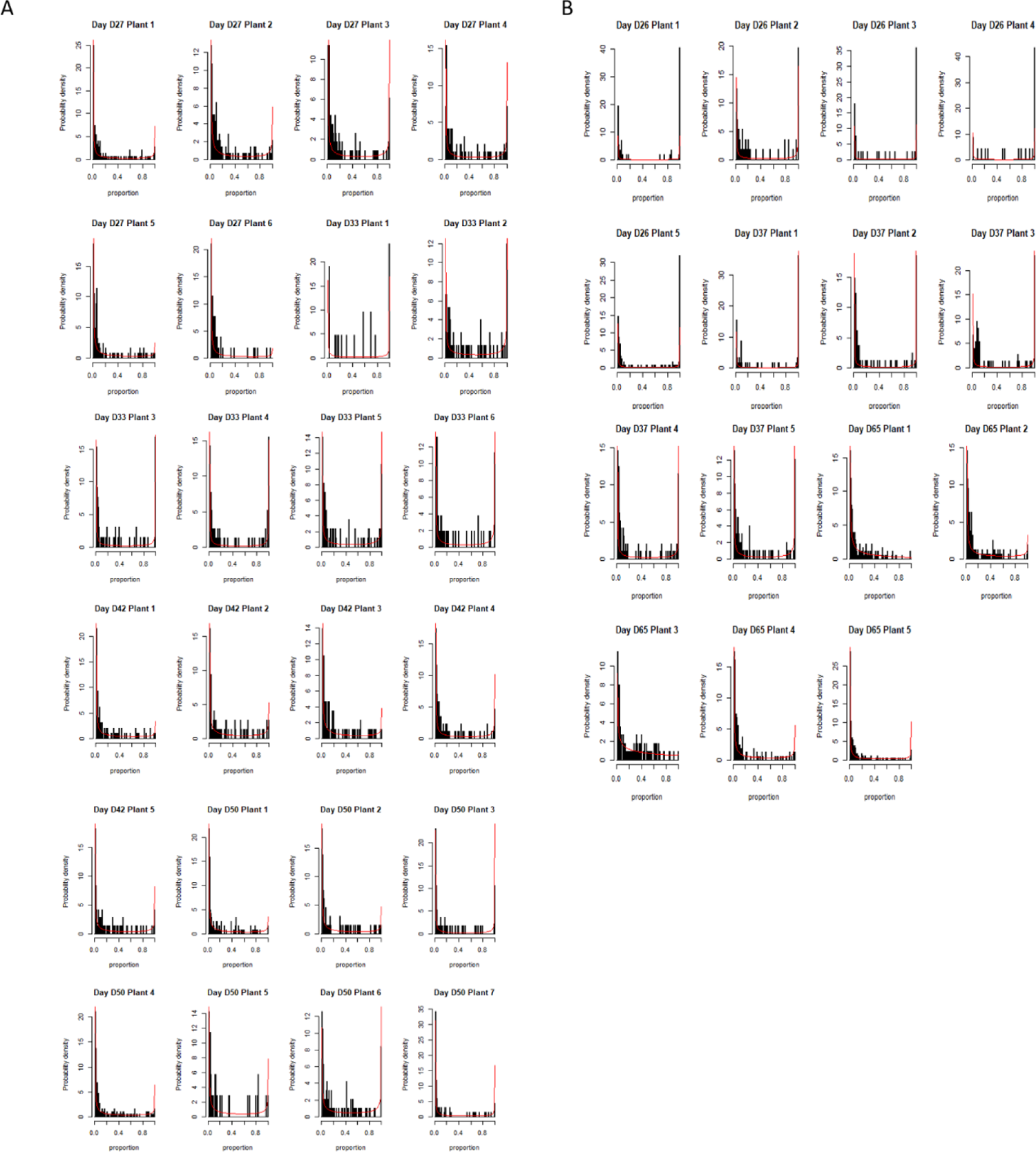
The empirical and estimated probability densities of proportions. Probabilities were estimated for each day and each plant of bean (**A**) and radish (**B**).

